# A neurophysiological alignment metric reveals a supportive role of unexpected stimuli in naturalistic language comprehension

**DOI:** 10.1101/2025.10.27.684752

**Authors:** Ashley L. M. Platt

## Abstract

Individuals use language every day to communicate their thoughts, emotions and expectations to those around them. Communication also incorporates receiving continuous and complex streams of information, not all of which is easily predictable. The memory literature suggests that this unexpected information may be processed more effectively when an individual possesses an adequate schema of the local context. We propose that, in language comprehension, we can determine the presence of a relevant scheme via an alignment metric, which compares an individual’s brain response to language (N400 amplitudes) with an expected response as determined by the probabilistic contingencies of the current context (surprisal). When alignment is higher, an individual’s internal model of the world is assumed to closely mirror the environment, constituting an effective schema. The present study sought to explore how individuals process unexpected information and whether successful comprehension is related to the alignment of internal predictive models to the external environment. It was hypothesised that higher alignment would support better comprehension of unexpected information than lower alignment. 40 participants (27F; mean age: 24.6 years [SD: 5.94]) listened to 12 short stories (3 genres, audio presentation) while their electroencephalogram (EEG) was recorded. Comprehension was tested via 6 multiple choice questions per story. A comprehension linear mixed-effects model (LMM) was computed for two regions of interest and two predictability metrics (N400 amplitudes and surprisal). A functional role of unexpectedness was demonstrated in all four models, where greater unexpectedness was related to higher comprehension scores. In one model we observed a significant interaction between alignment and average N400 amplitudes, indicating that comprehension was better for more unexpected stories when there was lower alignment to expected contextual probabilities. Our results suggest that individuals can utilise unexpected information to support language comprehension and that there are conditions where successful integration of this information is more likely. Contrary to our expectations, lower alignment to the current probabilistic conditions led to a greater capacity to utilise unexpected information. Future research should explore whether lower alignment reflects an individual’s openness to integrate new information into existing schemas and how this impacts comprehension of unexpected language.

## 1 Introduction

Language is a key component in everyday communication, providing individuals with the opportunity to share their internal thoughts with those around them. To support this functionality, language has infinite combinatoriality, thereby allowing individuals to communicate an infinite number of thoughts (e.g., Zuidema & de Boer, 2018). Consequently, individuals’ internal language processing systems are frequently faced with processing unexpected information. Studies which have explored the real-time processing of linguistic information have long suggested that less predictable words are associated with more processing ‘effort’, as reflected by increased reading and fixation times or more pronounced N400 event-related potential effects (e.g., Frank et al., 2013; Kutas & Federmeier, 2010; Smith & Levy, 2013). However, such studies are often highly constrained by design: for example, words are structured to be extremely predictable or unpredictable and target words are often placed towards the end of a sentence to allow for a clear comparison of the effect of predictability at a baseline processing level (Ferreira & Lowder, 2016). As such it is often concluded that, at least in these isolated processing contexts, increased predictability of language inputs is associated with successful comprehension and that therefore a primary goal of the language comprehension system is to reduce unexpectedness. To further consider the role of predictability from a communication perspective, these real-time processing studies have also been complemented by more naturalistic approaches, which encompass broader ranges of predictability. The present study seeks to add to these perspectives by drawing on conclusions from the memory domain and the account outlined in Platt (2025) which proposes how unexpected stimuli may support increased comprehension in certain processing contexts. In the following, we will outline how individuals utilise predictability during communication, consider the role of predictability outside of language processing and outline the account of *Functional Unexpectedness*.

### 1.1 Predictability and communication

Existing perspectives outline how language properties, such as predictability distribution and sentence structure, are key characteristics which can explain how information is communicated. Communication involves both the confirmation of existing knowledge and the sharing of new, potentially unexpected information. Without receiving or seeking out a wide range of predictable or unpredictable information, the language system would not have the capacity to reinforce existing probabilities or update knowledge in response to new information. Individuals would remain underdeveloped or overly flexible in their predictive models without this balanced input. However, often language research has focused on the importance of predictable information, over and above unpredictable. Receiving unknown or unexpected information may be what initially guides an interaction and it has been established that there is an inverse relationship between a word’s predictability and the degree of information it contains (Hale, 2016; Levy, 2008): from an information-theoretical perspective (i.e., surprisal; Shannon, 1948), linguistic units with higher surprisal, and therefore lower predictability, have a higher information load (Xu & Futrell, 2025). This supports the notion that unexpected inputs are essential to communication and highlights the importance of considering how unexpected, and potentially more informative, language is processed and understood.

An example of how language is shaped to support communication is that sentences do not uniformly distribute information; instead, already known information is placed first. This allows the comprehender to link the sentence to broader discourse, before then processing the new information. This perspective is outlined by Haviland & Clark (1974) in their Given-New strategy, which suggests that listeners attempt to identify given and new information in order to integrate new information into existing memory structures. Further, linguistic devices exist to support this structure, including discourse markers (e.g., “similarly” or “in contrast”) or expressions (e.g., “too”). Prosody also supports a given/new structure with focal stress often occurring on new information (Chafe, 1974, 1976; Halliday, 1967; Most & Saltz, 1979). Ferreira & Lowder (2016) suggest that structuring communication in these ways allows for dedication of processing resources to the integration of new information and that the role of given information is to highlight the existing structures that new information can be integrated into. These observations demonstrate how the language system is designed not only to handle unexpected information but also to utilise language structure to support communication goals. Successful comprehension is therefore not about having predictions precisely confirmed – in fact, perfect predictability is rare rather than typical in real life (Ferreira & Lowder, 2016; Luke & Christianson, 2016). Instead, successful comprehension is about integrating new information and the prediction-related mechanisms which support this task. Therefore, a communication-centred research approach might consider how individuals utilise unexpected inputs to acquire new information and should therefore seek to separate the role of predictability on real-time comprehension and the understanding of information for later use (i.e., during a follow up comprehension task). This might entail, for example, focussing language comprehension measures on how well individuals have understood the information they have processed rather than only how predictability influences real-time behavioural measures such as reading time.

### 1.2 How is unexpectedness useful?

Language is not the only domain of processing in which predictability plays a key role in determining the effectiveness of the system. In fact, it is often assumed that the brain acts as a ‘prediction machine’ and is constantly predicting incoming information across all domains of processing (Clark, 2013; Friston, 2008). In contrast to the majority of language comprehension research, other domains such as learning and memory research often operate on longer time frames and are therefore inherently designed to explore longer term effects of predictability. Perhaps as a key implication of this longer timeline, memory and learning research has demonstrated a supportive role of unexpectedness on behavioural outcomes such as memory recognition (Corley et al., 2007; Haeuser & Kray, 2023) or learning a new language (Lai et al., 2019). Additionally, memory research demonstrates that it is useful to remember both highly expected and highly unexpected events (Bein et al., 2023; Quent et al., 2022). One theory of these processes, the schema-linked interaction between medial prefrontal and medial temporal lobe model (SLIMM) proposed by van Kesteren et al. (2012) proposes that different brain systems support memory outcomes at these extremes, producing a U-shaped relationship between predictability and memory. The foundation of this framework resembles the Given-New structure outlined above, in that integration of information via the activation of certain brain networks is determined by how well stimuli fit into preexisting memory structures. These preexisting structures are commonly referred to as schemata and defined as activated world knowledge which is relevant to the current situation (Bein et al., 2023; Quent et al., 2022). For example, when considering the neocortical representation of items in a kitchen, items which are frequently associated with a kitchen schema (e.g., toaster, fridge or microwave) are posited to activate the medial prefrontal cortex (mPFC) and inhibit the medial temporal lobe (MTL) (van Kesteren et al., 2012). When an unpredictable item is perceived in the kitchen, (e.g., a couch or pillow), a lack of association between the activated representations (i.e., kitchen and couch) results in inhibition of the mPFC and activation of the MTL. In a memory task, both conditions would result in improved memory for these items in comparison to a more neutral item (e.g., broom). Evidence for this U-shape relationship was demonstrated by Quent et al. (2022) who showed that participants were able to more accurately recall items which were highly expected or highly unexpected in a specific environment.

Complementary to findings in the memory domain, there is evidence that errors in prediction can promote memory encoding during language processing. Termed error-driven learning, this is a process by which prediction errors drive the formation of new memories (e.g., Kalbe & Schwabe, 2022; Loock et al., 2025). This perspective suggests that by engaging in experiences which increase prediction errors, individuals are provided with a better opportunity to support learning. Unlike more traditional language-based studies, this approach aligns with the notion that unexpectedness is useful in information gathering and preparing the prediction system for future unpredictable events (Parr et al., 2022). For example, Hodapp & Rabovsky (2021) examined N400 amplitudes within a sentence reading task as a measure of prediction error and observed faster reaction times in an implicit memory task for words that had previously elicited larger N400 effects in the preceding reading task. Based on these results, they argue that the N400 can be viewed as an implicit error-driven learning signal (Hodapp & Rabovsky, 2021). Haeuser & Kray (2023) also demonstrate the usefulness of prediction errors, concluding that less expected nouns (as indexed by larger N400 amplitudes) in a comprehension task are more readily identified in a follow-up word recognition task. Additionally, a frequent example of how prediction errors support learning is in the language acquisition of young children. Children as young as 9 months show prediction errors (larger N400 amplitudes) when presented with pictures that are incongruent to spoken words (Parise & Csibra, 2012). It is commonly concluded that children use these prediction errors to update models and support the further development of language (Kray et al., 2024; Zettersten, 2019). These studies demonstrate that there is a key role of prediction errors in improving behavioural outcomes across multiple domains of processing. Drawing on these perspectives, the following section will outline the account of unexpected information is processed with respect to comprehension depth (i.e., functional unexpectedness) and suggestions for how to explore this.

### 1.3 Functional Unexpectedness

In Platt (2025), a framework was proposed outlining how to measure individual alignment to local linguistic context and how this alignment may indicate a functional role of unexpectedness during language comprehension. It was suggested that unexpectedness is useful for information processing when an individual possesses suitable predictive schemas which support the integration of new, unexpected information. This perspective built on conclusions from SLIMM (van Kesteren et al., 2012), a model which highlights that memory for unexpected items is improved only when integration with preexisting memory structures is possible. This account will be empirically tested in the present study by including the following three key components:

1. A measure of how expected language input is given the current context
2. A measure of the extent to which the language user possesses a suitable schema for the context in which the current input item occurs
3. A suitable outcome measure for assessing the effects of unexpectedness Prior linguistic research has determined that the predictability of language input in the current context can reliably be measured by corpus-based metrics such as surprisal. Surprisal is a context-based metric of predictability: the surprisal value of a word w_i_ is defined as the negative log probability of w_i_ given the previous words in a sentence, or a previous word window of any length (Hale, 2006):

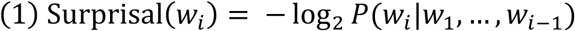

Surprisal theory (Hale, 2006; Levy, 2008, for a review see Hale, 2016) builds on the foundation that comprehenders use probabilistic knowledge from past linguistic experience, perhaps akin to the schemas relied on in memory, to generate expectations about current and upcoming information. As first demonstrated by reading time studies, increased surprisal has been tied to greater processing difficulty (Lowder et al., 2018; Smith & Levy, 2013; Venhuizen et al., 2019). Subsequently, advances in predictive language models now allow the calculation of surprisal from Large Language Models (LLMs) which are trained on billions of words and have the capacity to incorporate all previous context (i.e., all previous words in a story). This allows surprisal values to more closely reflect the linguistic experiences of a comprehender, who has access to an entire lifetime of linguistic experience. In the present study, surprisal measures will be incorporated in an alignment metric, outlined in more detail in the following, and as a key predictor of comprehension depth.

In order to address the second component of functional unexpectedness, Platt (2025), suggests that the relationship between corpus-based surprisal and N400 amplitudes can provide a metric of individual alignment. Unlike in memory and learning tasks, it is harder to ascertain an individual’s unique level of linguistic experience without jeopardising the applicability of findings to a real-life context (i.e., by providing extremely predictable continuations within highly constrained sentences). This alignment metric will seek to operationalise the presence of a predictive model or schema at an individual and story level. With word-by-word surprisal acting as a ‘baseline’ measure of predictability, N400 amplitudes can measure an individual’s tailored experience of the linguistic stimuli. The role of surprisal in this alignment metric requires calculations to include a large contextual window these calculations are proposed best represent expected responses to linguistic stimuli based on the probabilities of language and the current context. The N400 component is a (typically) centro-parietally distributed, negative-going ERP component which peaks between 300-500ms post word onset (Kutas & Hillyard, 1980). Its amplitude is negatively correlated with stimulus predictability, with less predictable stimuli eliciting larger (more negative) amplitudes compared to predictable stimuli. Larger N400 amplitudes in response to linguistic stimuli have been associated with increased text difficulty and reading times (Szewczyk & Schriefers, 2018). Previous research has also indicated that surprisal correlates with N400 amplitudes (Aurnhammer & Frank, 2019; Frank et al., 2013). In view of this relationship, it is often posited that the N400 reflects predictive processes in language (e.g., Bornkessel-Schlesewsky & Schlesewsky, 2019; Frank et al., 2015; Kuperberg & Jaeger, 2016; Rabovsky & McRae, 2014) and that language comprehension operates most effectively in conditions of high predictability, as indexed by smaller N400 responses (Dambacher et al., 2006; DeLong & Kutas, 2020; Payne et al., 2015; Roehm et al., 2007). Platt (2025) suggests that the more aligned an individual’s word-by-word N400 amplitudes are with surprisal, the more aligned their predictions are to local context. Further, it is proposed that individuals with higher alignment can suitably integrate new, unexpected information into existing predictive schemas for use in future communicative contexts. In the present study we predict that individuals with higher alignment will have greater comprehension depth than those who are less aligned to local context.

Finally, to measure an individual’s contextual understanding of naturalistic stimuli, the present study has designed a series of comprehension questions to follow a story listening task. These questions were designed to probe the meaning individuals have gathered from the linguistic information and therefore provide an indication of their comprehension depth. A measure of comprehension depth is proposed to be situated between traditional memory literature measures (e.g., recall or recognition memory tests) and follow-up metrics commonly explored in language comprehension studies (e.g., reading and fixation times or comprehension probes designed only to measure whether a listener is paying attention). This ensures an outcome measure which probes depth of contextual understanding. Consideration for a depth of comprehension metric in addition to an alignment metric has the potential to resolve the intriguing discrepancy in that language comprehension settings which elicit larger N400 amplitudes are framed as less favourable for processing while simultaneously being associated with downstream advantages such as improved memory and learning.

### 1.4 The current study

The present study sought to explore the role of predictability in naturalistic language comprehension by testing the predictions made in Platt (2025). This requires the inclusion of individual alignment and depth of comprehension metrics to examine a possible functional role of unexpectedness. Participants listened to short stories of different genres while their electroencephalogram (EEG) was recorded, with each story followed by a series of questions designed to probe depth of comprehension. Individual levels of alignment, as determined by the alignment of individual-level N400 amplitudes and surprisal (with a context window of the entire story), were utilised to predict depth of comprehension outcomes. Additionally, these comprehension models included either story-average N400 amplitudes or surprisal as a key predictor, to provide an additional comparison of a corpus-based versus an individual-level metric of predictability. Separating these predictability metrics into separate models provided insight into how the novel relationships (i.e., alignment and depth of comprehension) differed when considering two common measures of predictability. It was hypothesised that a functional role of unexpectedness would be present under conditions of higher alignment, in that lower predictability would be associated with greater comprehension depth.

## 2 Methods

### 2.1 Participants

Forty participants (27 female) participated in the current study. All were healthy, right-handed adults between the ages of 18 and 40 years (mean: 24.6, SD: 5.94). Handedness was assessed using the Flinders Handedness Survey (Nicholls et al., 2013), a brief self-report measure of skilled handedness. They had not taken recreational drugs in the last 6 months and were not taking regular medication which could affect the EEG. Participants also had normal or corrected vision, reported normal auditory acuity and did not have any diagnosed psychiatric, neurological, cognitive or language disorders. Additionally, to avoid language confounds, participants were native English speakers who had not learned a second language before the age of 5 years. The experiment protocol was approved by the University of South Australia’s Human Research Ethics Committee (protocol number: 204945).

### 2.2 Materials and measures

#### Audio stimuli

The audio stimuli for this experiment included 12 short stories classified as fiction, newspaper and science (4 stories per genre) with an approximate duration of 3-4 minutes. Stories were recorded in mono format (sampling rate: 44kHz) by a native speaker of Australian English. A number of characteristics per story were calculated for use in the data analysis. This included word position, word duration, sentence length, word frequency, idea density, word audio frequency and intensity. WebMAUS Basic (Schiel, 1999) was used to calculate word onsets and offsets. Unigram frequency values were computed from the Open Subtitles corpus for English (751 million words) as made available by van Paridon & Thompson (2021). Computerised Propositional Idea Density Rater (CPIDR; Brown et al., 2008) was used to calculate ID for each short story. Idea density (ID; also known as Propositional Density or P-Density. Kintsch & Keenan, 1973) uses written or oral text samples to calculate the number of ideas expressed relative to the number of words used and has been linked to linguistic ability (Farias et al., 2012), language production (Kemper et al., 2001) and comprehension (Kintsch & Keenan, 1973). Acoustic characteristics were calculated using Praat version 6.4.04 (Boersma & Weenink, 2024). The current study used this software to calculate word duration, fundamental frequency and intensity. The descriptive characteristics of each story are outlined in Table 1. See Supplementary Material A for story stimuli examples.

**Table 1:** Characteristics for audio stimuli at genre level [M (SD)].

| Genre | Mean duration | Mean word count | Mean Idea density | Mean Word frequency | Mean Word duration | Mean Sentence length | Mean Fundamental frequency | Mean Intensity | Mean Surprisal |
| --- | --- | --- | --- | --- | --- | --- | --- | --- | --- |
| Fiction | 2m50s | 430.0<br>(42.1) | 0.52<br>(0.03) | 5014689.5<br>(589760.1) | 0.30<br>(0.03) | 14.41<br>(5.51) | 227.6<br>(6.96) | 58.53<br>(0.55) | 5.30<br>(0.77) |
| Newspaper | 2m57s | 433.5<br>(11.5) | 0.46<br>(0.04) | 5161020.6<br>(707570.0) | 0.33<br>(0.01) | 26.30<br>(3.43) | 227.9<br>(1.84) | 57.88<br>(0.54) | 5.73<br>(0.65) |
| Science | 3m10s | 446.8<br>(57.8) | 0.52<br>(0.05) | 4465183.5<br>(637829.1) | 0.35<br>(0.01) | 19.85<br>(1.74) | 225.6<br>(5.86) | 57.29<br>(1.44) | 5.17<br>(0.24) |

#### Comprehension probes

Depth of comprehension was measured using story-relevant multiple-choice questions presented at the end of each story. Six questions were presented for each story, resulting in a total of 72 across the experiment. Questions were designed based on an approach outlined in Delgado & Salmerón (2021) and Ozuru et al. (2007). These studies suggest that comprehension questions can be designed to probe a contextual understanding of stimuli and not only rely on recall memory. As such, the final questions were chosen to reflect a listener’s in-depth contextual understanding of the stories, beyond the surface level information provided by text. Comprehension probes were piloted for clarity and to ensure ceiling effects were not present.

Depth of comprehension for modelling, as indicated by comprehension probes, was calculated at story level to reflect the number proportion of correct answers (i.e., a score ranging from 0-1). See Figure 1 for example of comprehension questions.

**Figure 1:**
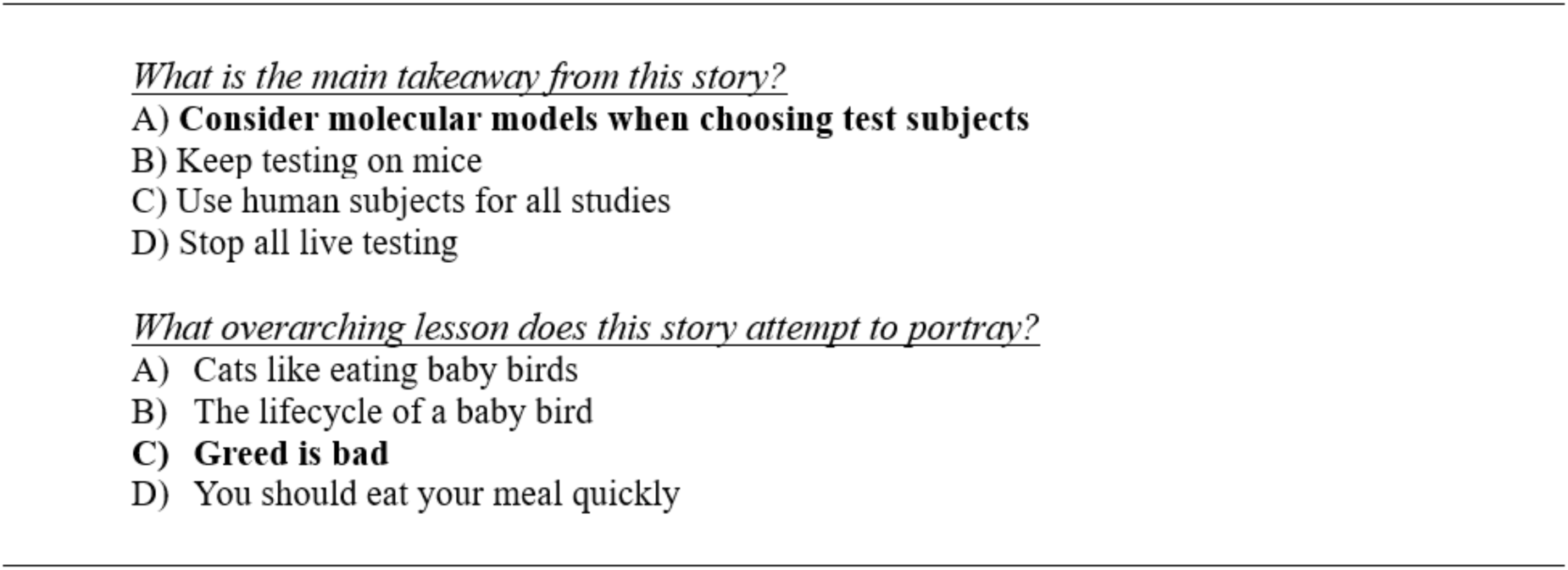
Example of multiple-choice comprehension depth questions.

#### Surprisal calculation

Surprisal was calculated at word level and included a context window of all words prior to the current word. Thus, the amount of contextual information used for the surprisal calculation increased over the course of each story. Word-level surprisal values were used in the alignment models and were then averaged at story-level for the comprehension models. Surprisal is the negative log probability of a word given its preceding context (Lowder et al., 2018) (see Equation 1).

To increase the context window of the surprisal calculations GPT-2 (Generative Pre-trained Transformer-2) was used, which is a pretrained large-scale transformer model trained on billons of words of text with the objective to predict the next word (Radford et al., 2018, 2019). GPT-2 was utilised via functions *GPTConfig, GPT2Tokenizer, GPT2LMHeadModel and GPT2Model* from the *transformers* package in Python version 3.10.14. Occasionally GPT-2 calculations split up words into ‘tokens’. In these cases, we summed the value of the tokens to provide the whole word surprisal, following Stanojević et al. (2023).

#### Electroencephalography (EEG) recording and pre-processing

EEG was recorded with a 32-channel Brainvision ActiCHamp system (Brain Products GmbH, Gilching, Germany) with a sampling rate of 500 Hz. Electrodes were placed according to the 10-20 system. Vertical and horizontal eye movements were monitored with two electrooculogram (EOG) electrodes placed beside the outer canthus of the right and left eyes, as well as two additional EOG electrodes slightly above and below the left eye. The online reference and ground were AFz and FPz, respectively. Impedances were kept below 10k Ω. EEG pre-processing and analysis was carried out in MNE Python version 1.6.1 (Gramfort et al., 2013). EEG signals were re-referenced to the average of left and right mastoids (TP9 & TP10). Independent Component Analysis (ICA) was utilised to correct EOG artifacts. Independent components found to correlate most strongly with EOG events (via the *create_eog_epochs* function in MNE) were excluded. A band-pass filter of 0.3-20Hz was applied to the raw data to remove signal drift and high frequency noise. The following analyses are based on epochs from −0.2 to 1 second relative to the onset of each word in each story.

### 2.3 Procedure

Interested participants completed a demographic questionnaire to ensure they fit study criteria. If eligible, participants were invited to attend a 2 hour in-lab testing session which included EEG setup and testing in addition to collection of linguistic and neurophysiological metrics not utilised in this analysis, such as resting state EEG and individual-level idea density. The story task ran for roughly 50min and required participants to listen to each story and then answer the six associated comprehension questions. Participants were initially presented with two instruction screens and prompted to ‘press any key to begin’. Each story was preceded by a 10 second count down before presenting a fixation cross at story onset and for the duration of each story. When the story was complete, further instructions were provided to press any key to continue to the comprehension questions. Participants were also prompted to take this time to relax before continuing with the experiment. The story-question interim instruction screen and comprehension questions were not given a maximal reaction time, and the experiment would only progress following participant engagement. Audio recordings were played through a loudspeaker at a comfortable volume for the participant. Story order presentation was determined by assignment to one of four pseudo-randomly generated lists, in which stories of the same genre never appeared more than twice in a row. List presentation was counterbalanced across participants.

### 2.4 Data analysis

Data analysis was approached in three-stages (see Figure 2 for overview). Stage 1 utilised a whole-head single trial analysis to identify tempo-spatial regions of interest (ROIs) in which surprisal and EEG amplitudes were significantly correlated. An identified ROI and a traditional N400 ROI were then used to extract ERP amplitudes for stage 2. Stage 2 computed a surprisal model per ROI to produce an alignment value per participant per story. These alignment values were then utilised in Stage 3 to compute a depth of comprehension model per ROI (whole-head analysis and traditional N400) and predictor (surprisal and average EEG amplitudes).

**Figure 2:**
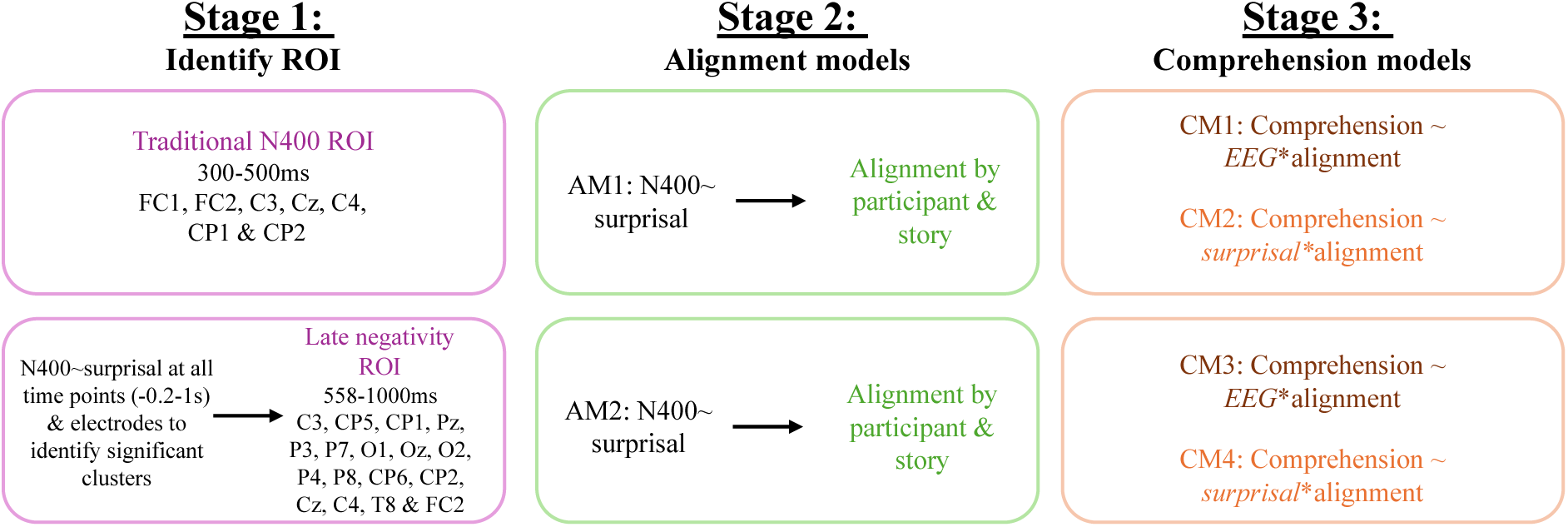
Overview of analysis pipeline.

#### Stage 1: Whole-head single trial analysis

Following Brennan & Hale (2019), a whole-head single trial analysis was carried out per-participant and across each time-point (−0.2-1s) and electrode to identify spatio-temporal clusters of EEG activity which correlate with surprisal. To this end, we calculated regression-based ERPs (rERPs; Smith & Kutas, 2015) for all content (i.e., adjectives, nouns, verbs & adverbs) words in each story. Parts of speech tagging was facilitated by the *nltk* package in Python and included the functions *word_tokeniser* and *pos_tag*. A regression model was calculated for each sample point and electrode, thereby allowing for the examination of predictors of interest while controlling for additional categorical and continuous covariates of no interest. This type of analysis is particularly well suited to naturalistic stimuli. In addition to surprisal as our primary predictor of interest, control predictors for the rERP analysis included pre-stimulus activity (−200-0 ms prior to word onset; Alday, 2019), word-position in the story, word duration, log-transformed word frequency, average fundamental frequency and intensity. Log-transformed word frequency (previous and following word) and previous word duration were also included as control predictors. Null-regression models, created by randomly permuting the rows of the design matrix, were then used in a group-level analysis. This group-analysis compared the beta coefficients for each effect to be tested across time points (0-1s) and across all electrodes using a non-parametric permutation test (Maris & Oostenveld, 2007). An F-test was conducted per time point and electrode, comparing target beta values with matched values from the null model. Significant (p <.01) electrode and timepoints were clustered based on spatio-temporal adjacency. These steps were repeated for 1000 permutations and significant clusters were identified (α = 0.05).

#### Stage 2: Alignment models

Individual ERP N400 amplitudes were calculated as the average voltage post word onset using regions of interest (ROI) as identified by the whole-head analysis and also from a more traditional N400 ROI across 300-500ms. The modality of linguistic stimuli has been shown to influence the topography of resulting ERP components (Holcomb et al., 1992; Holcomb & Neville, 1990). In particular, auditory N400 effects have demonstrated a broader distribution that the centro-parietal distribution elicited by visual stimuli (Wolff et al., 2008). As such, the present study calculated N400 amplitudes from the following electrodes: FC1, FC2, C3, Cz, C4, CP1 and CP2. EEG data were analysed using linear mixed effects models (LMMs) with the MixedModels.jl package in Julia (Bates et al., 2021). These models included surprisal, log unigram frequency, word position and prestimulus activity and their interactions as fixed effects. All control predictors that were used in the rERP analysis were also included., word duration (previous & current), log-transformed previous word frequency, average fundamental frequency and intensity. The random effects structure for each model was determined using a parsimonious model selection procedure to avoid over-parameterisation (Bates et al., 2021; Matuschek et al., 2017), with final models including random intercepts by subject, story, word and channel. Final model structure is outlined below and was consistent between the two ROIs.

***Alignment model structure:*** average n400 amplitudes ∼ pre-stimulus activity * word position * log frequency * story-level surprisal + duration + average fundamental frequency + average intensity + previous word frequency + previous word duration + (1 + pre-stimulus activity + log frequency + word position | subj) + (1 + story-level surprisal | subject & story) + (1 + pre-stimulus activity + word position + story-level surprisal | word) + (1 | channel)

#### Stage 3: Comprehension analysis

The above alignment model (per window) was utilised to compute the random slope of surprisal by participant for each story. As a proxy for individual-level alignment to story context, these beta values were then used to predict comprehension outcomes. For each window, two models were computed, one including a story-average EEG predictor and the second a story-average surprisal predictor. Each of these models also include a quadratic term for surprisal and N400 amplitude predictors. Inclusion of a quadratic term allowed the model to account for non-linear relationships, as theorised in Platt (2025). Each model included story level control predictors of average sentence length, average word duration, average log transformed word frequency, idea density and genre. Random effects structures were selected using the same procedure as stage 2 with final random effects structures are outlined below in Table 2.

**Table 2:** Random effects structures for all comprehension models.

| Window | Model | Random effects structure |
| --- | --- | --- |
| Traditional N400 | EEG | Idea density \| subject<br>1 \| story |
|  | Surprisal | Average sentence length \| subject<br>1 \| story |
| Late Negativity | EEG | Average sentence length \| subject<br>1 \| story |
|  | Surprisal | Average sentence length \| subject<br>1 \| story |

*Note: control predictors were consistent across all models*

## 4 Results

### 4.1 Behavioural outcomes

Participant comprehension scores are outlined in Table 3. The average participant comprehension score across genres was 70.0%. Two participants scored below 50% for the science genre, however their mean score across genres was above 50% and therefore were not removed from analysis.

**Table 3:** Summary of participant comprehension scores.

| Genre | Total questions (N) | Mean % (SD) |
| --- | --- | --- |
| All Stories | 72 | 70.0 (9.48) |
| Fiction | 24 | 72.9 (11.3) |
| Newspaper | 24 | 69.7 (13.9) |
| Science | 24 | 67.4 (11.0) |

### 4.2 Frequency effect ERP

As a sanity check for our naturalistic design, we computed EEG grand averages by word frequency. The N400 component has been shown to be modulated by frequency with higher N400 amplitudes for low versus high frequency words (Kutas & Federmeier, 2011) and as such it is suitable to assume that this effect will be present in our analysis. Figure 3 illustrates the effect of frequency and reflects peak N400 amplitudes at roughly 700ms post word onset. Although the latency of this effect is unexpected it is comparable to a previous study by (Alday et al., 2017) which explored the processing of naturalistic auditory stimuli and also demonstrated a frequency effect at around 700ms, attributing this latency to variations in word length, a characteristic which is not controlled in naturalistic stimuli.

**Figure 3:**
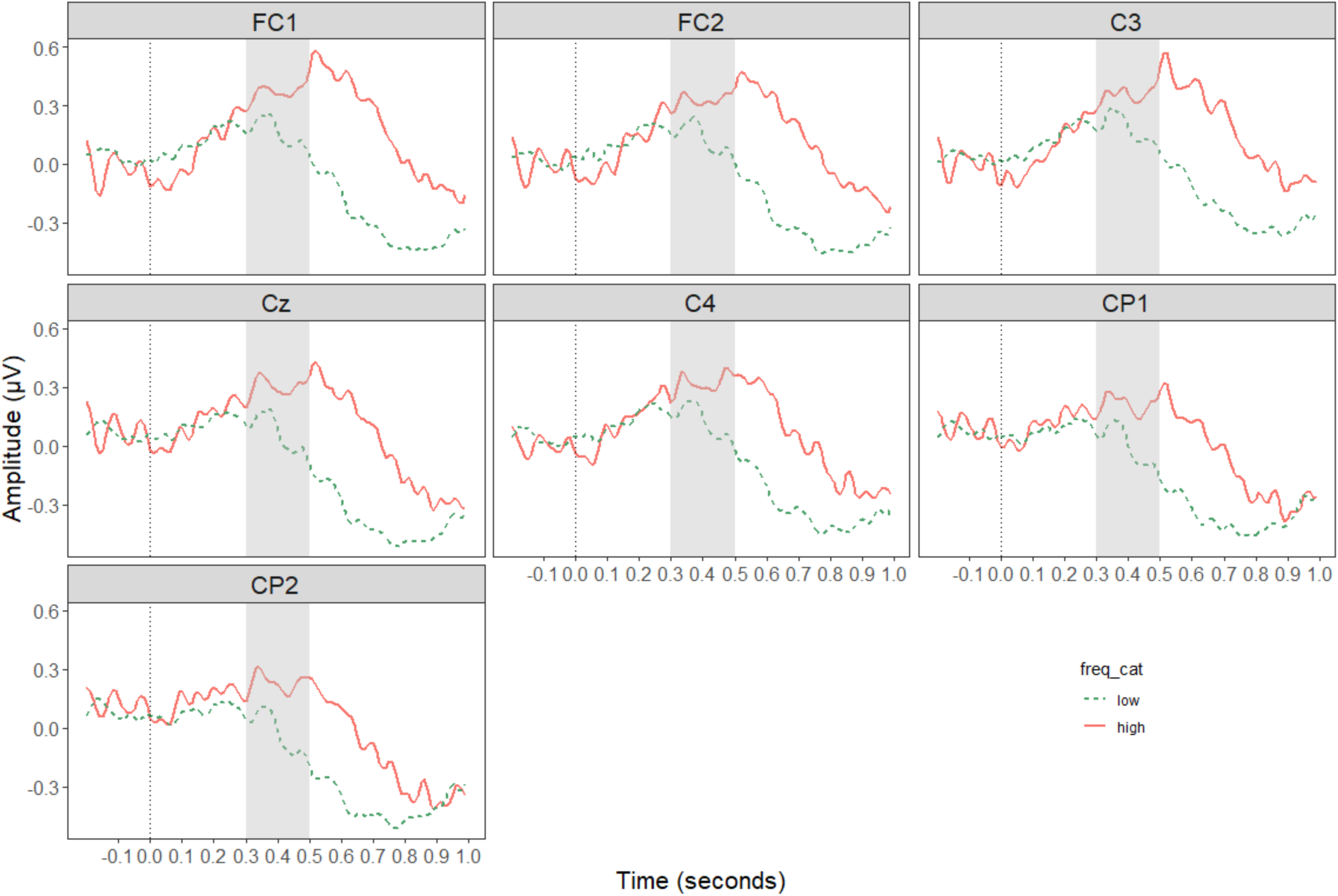
Grand average ERP plots of frequency effect. Lines are separated by word frequency and amplitudes are plotted with negativity downwards. Highlighted region identified N400 time window 300-500ms. Note: only content words are included in averages.

### 4.3 Stage 1: Whole-head single trial analysis

A spatio-temporal cluster was identified in the whole-head single trial analysis. This cluster ranged from 558-1000ms and encompassed electrodes: C3, CP5, CP1, Pz, P3, P7, O1, Oz, O2, P4, P8, CP6, CP2, Cz, C4, T8 and FC2. These characteristics were then utilised to compute average EEG amplitudes within this ROI for the alignment models. See Figure 4 for spatio-temporal characteristics of the cluster.

**Figure 4:**
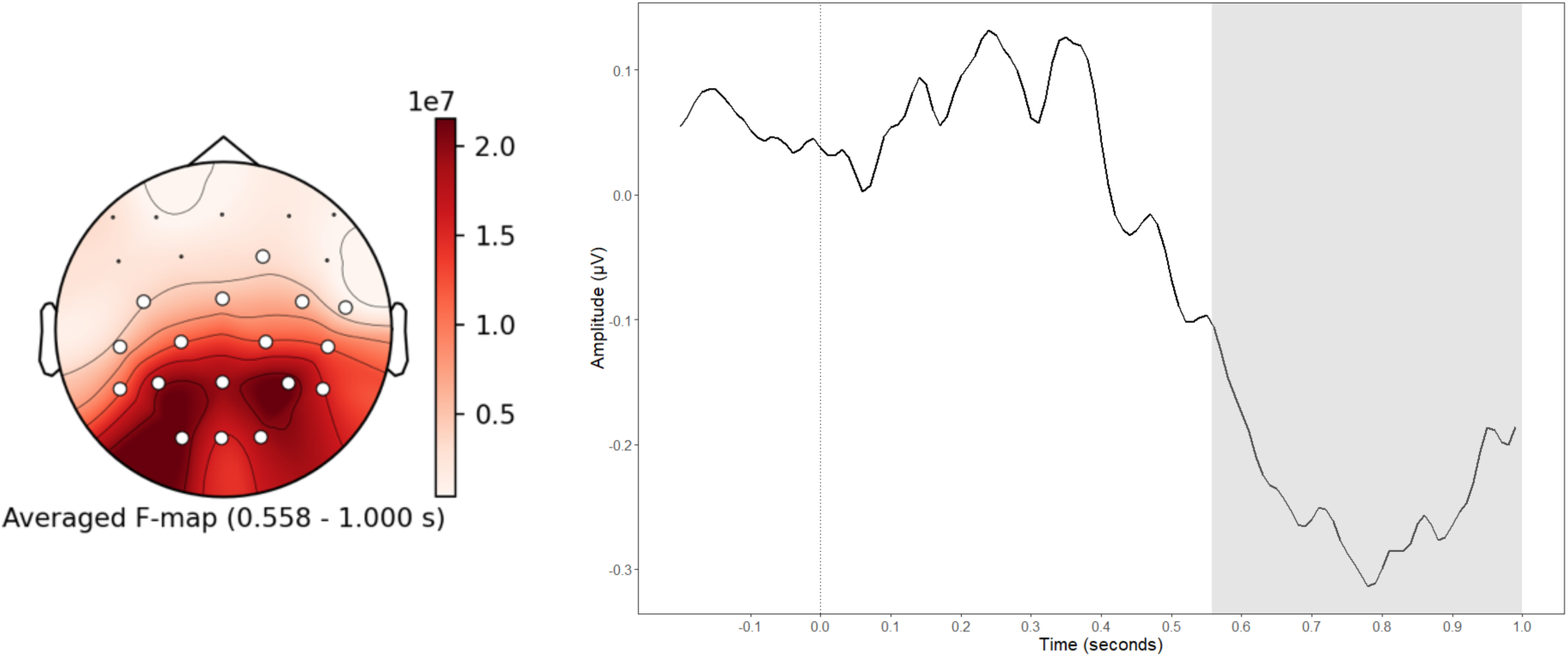
Spatio-temporal characteristics and grand average plot for audio cluster.

### 4.4 Stage 2: Alignment models

A mixed effects model was computed for each region of interest, traditional N400 (AM1) and late-negativity (AM2) using story-level surprisal. The top-level interactions between word position, log transformed word frequency and story-level surprisal were significant for both AM1 (Estimate = −0.06, Std. Error = 0.02, z = −3.43, *p* <.001) and AM2 (Estimate = −0.10, Std. Error = 0.01, z = −9.61, *p* <.001). This interaction effect is visualised in Figure 5 for AM2. See Supplementary Material B for model outputs.

**Figure 5:**
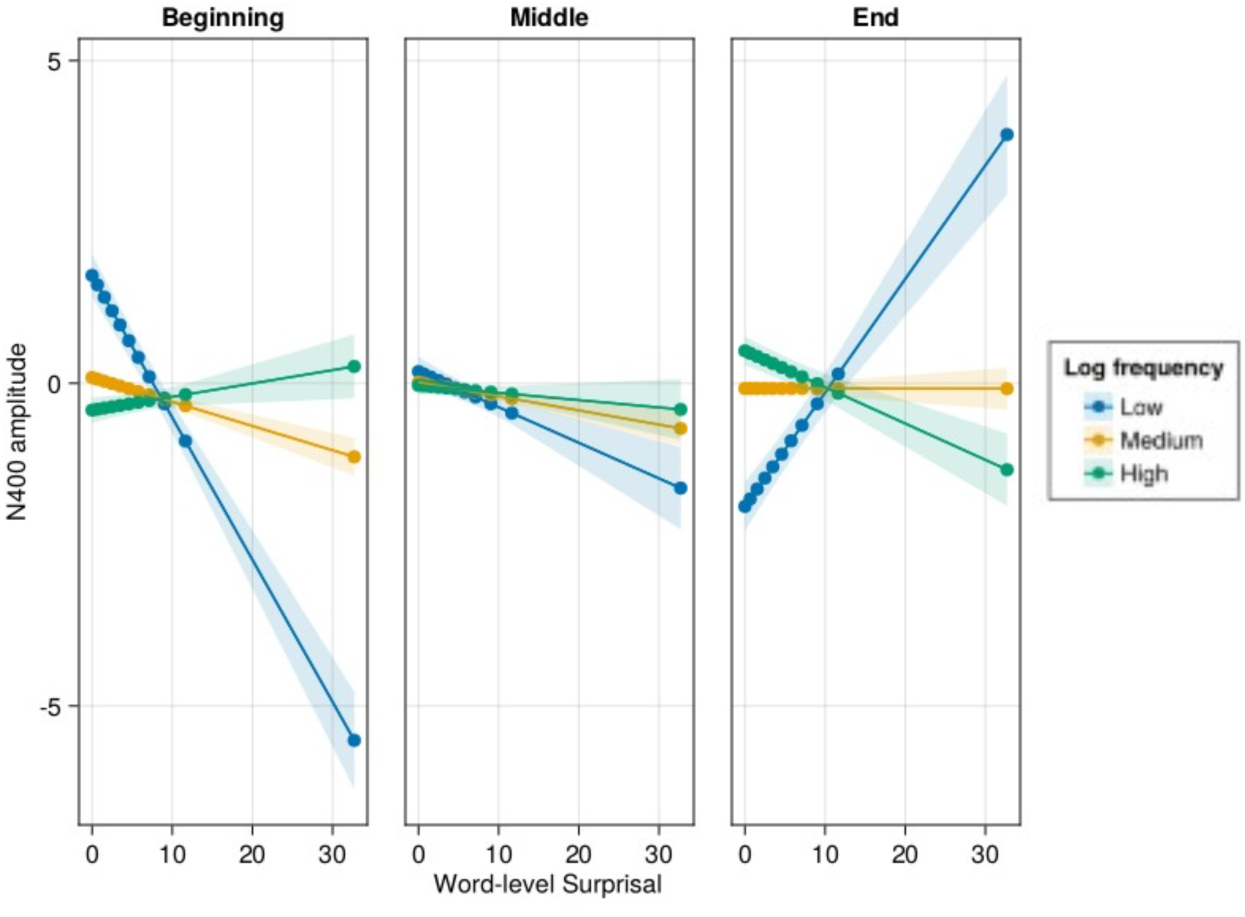
Interaction between N400 amplitudes and story-level surprisal by word frequency and position in story. The x-axis is surprisal, whilst the y-axis represents fitted values of N400 amplitude. The graph is faceted by story position and low, medium, and high frequency words are represented in separate slopes across each facet. The trichotomisation of word position and word frequency are for visualisation purposes only and were continuous predictors in the mixed effects model. Filled colours are 95% confidence intervals.

The random slopes of story-level surprisal by story and participant are utilised in the following comprehension models. The distribution of these values for each ROI is displayed in Figure 6. The varied distribution of these values suggests sufficient variability to utilise our alignment metric as an individual-level predictor of comprehension depth. Figure 7 visualises alignment per individual and highlights both individual and genre variability.

**Figure 6:**
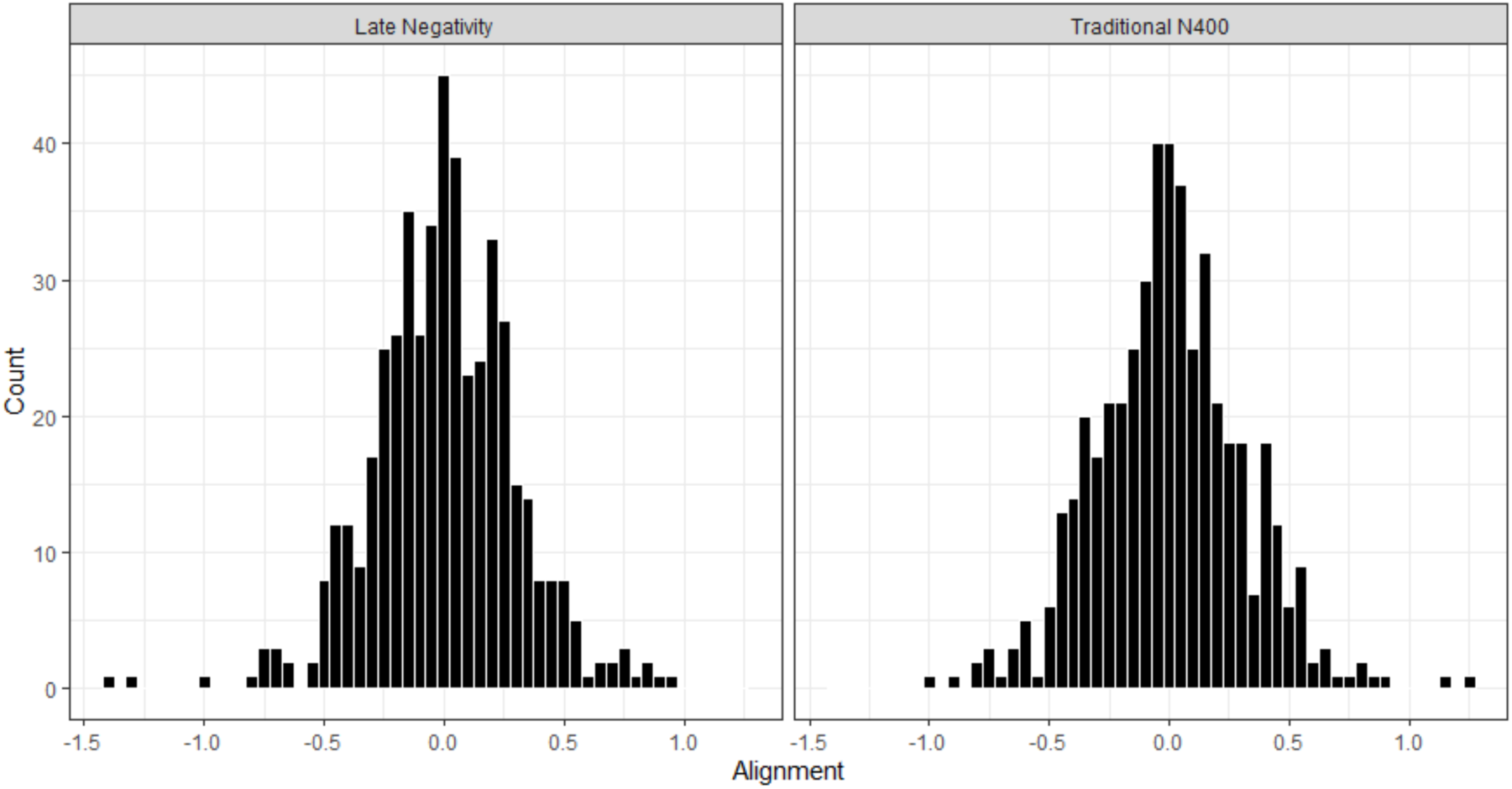
Distribution of alignment values for Traditional N400 ROI (LHS) and Whole-head analysis ROI (RHS).

**Figure 7:**
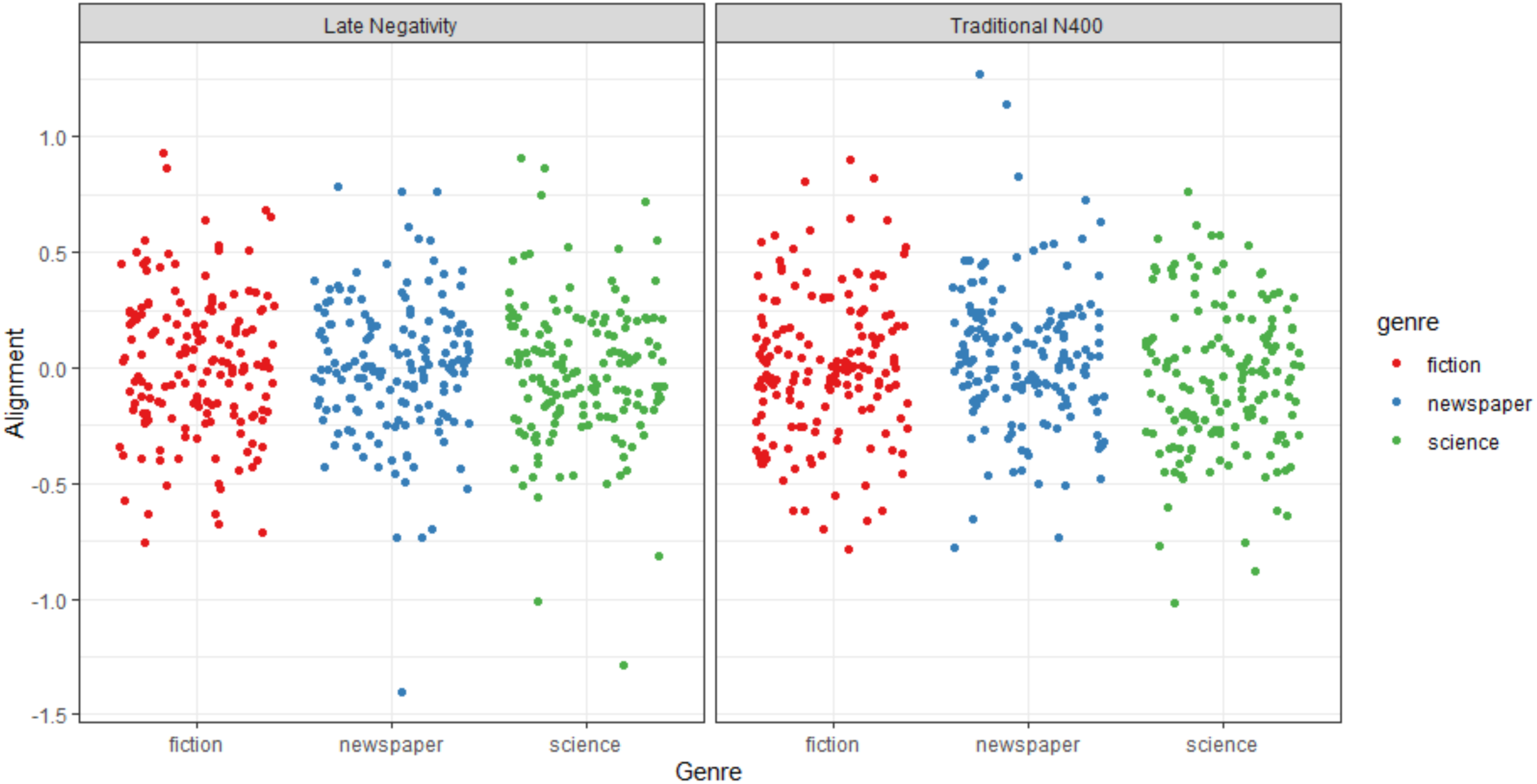
Individual variation or alignment values by genre, for Traditional N400 ROI (LHS) and Whole-head analysis ROI (RHS).

### 4.5 Stage 3: Comprehension models

A mixed effects model was computed for average EEG amplitudes (within each ROI) per story and average story-level surprisal per story for each region of interest. All models showed main effects of control variables, average sentence length, average word duration, idea density and genre. The surprisal models further reported an effect of average word frequency. See Supplementary Material B for full model outputs.

#### CM1: Traditional N400 ROI & EEG

This model showed a significant interaction effect between alignment values, average EEG amplitude and average EEG amplitude^2^ (Estimate = −0.02, Std. Error = 0.01, z = −2.47, *p* = .012). 2D and 3D representations of this interaction effect are visualised in Figure 8 and Figure 11 and illustrate that comprehension scores are highest at low alignment and more negative EEG amplitudes.

**Figure 8:**
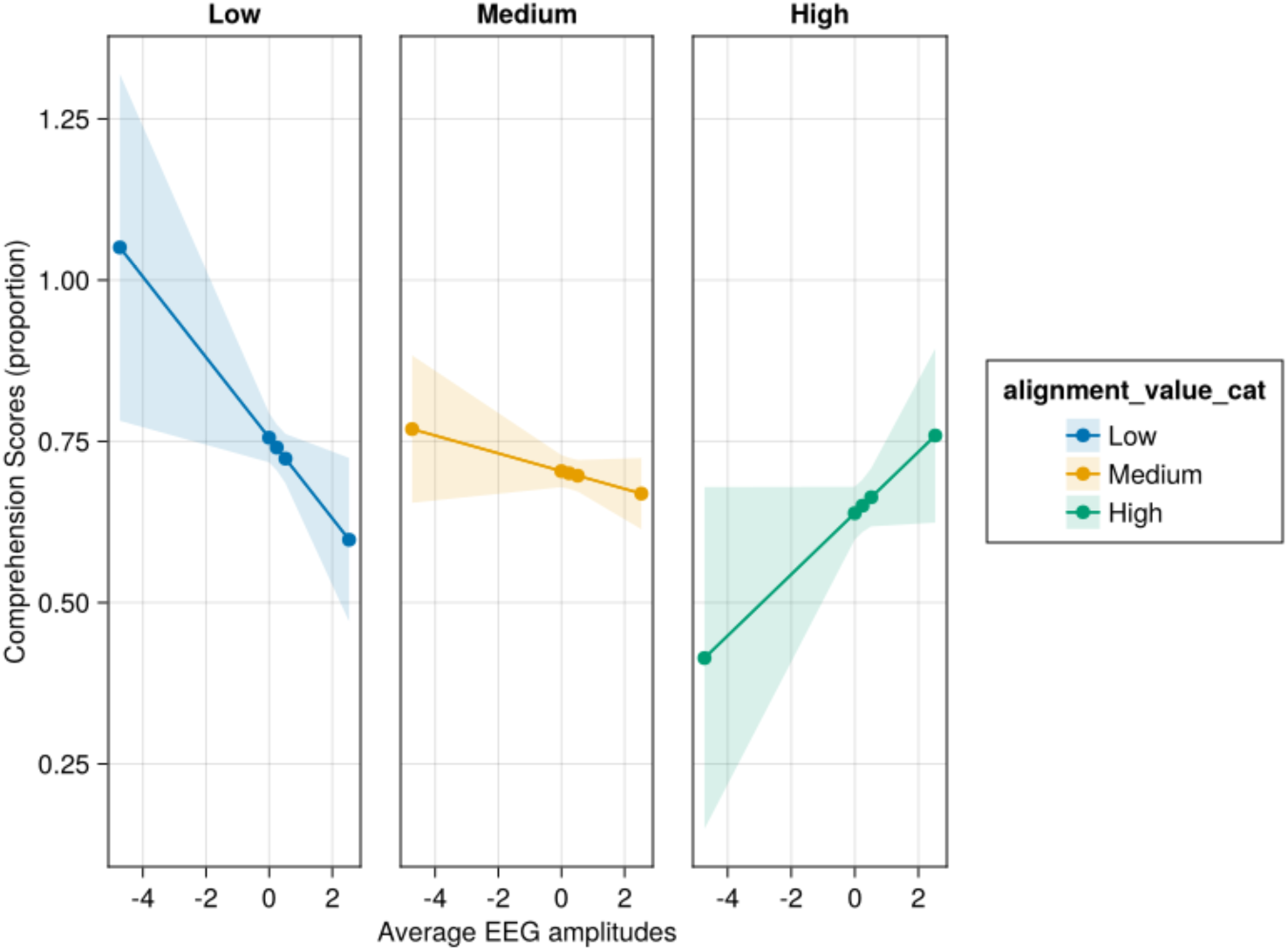
2D visualisation of the highest order effect (alignment*EEG amplitudes*EEG amplitudes2) for CM1. The x-axis is average EEG amplitudes, whilst the y-axis represents fitted values of comprehension scores. The graph is faceted by alignment and low, medium, and high alignment scores are represented in separate slopes across each facet. The trichotomisation of alignment is for visualisation purposes only and was a continuous predictor in the mixed effects model. Filled colours are 95% confidence intervals.

**Figure 9:**
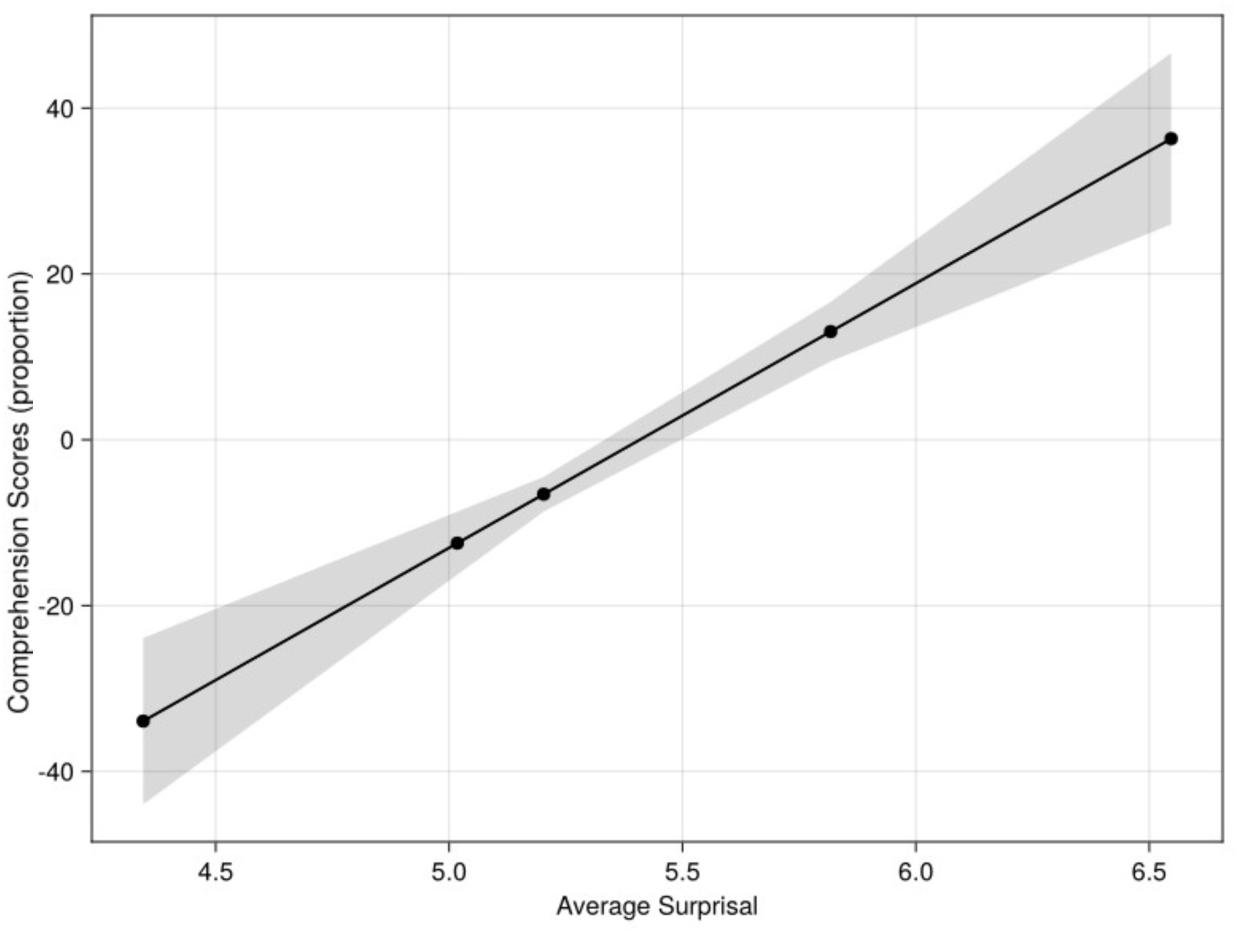
Main effect of average surprisal in CM2. The x-axis is average surprisal, whilst the y-axis represents fitted values of comprehension scores. The shaded areas represent 95% confidence intervals.

**Figure 10:**
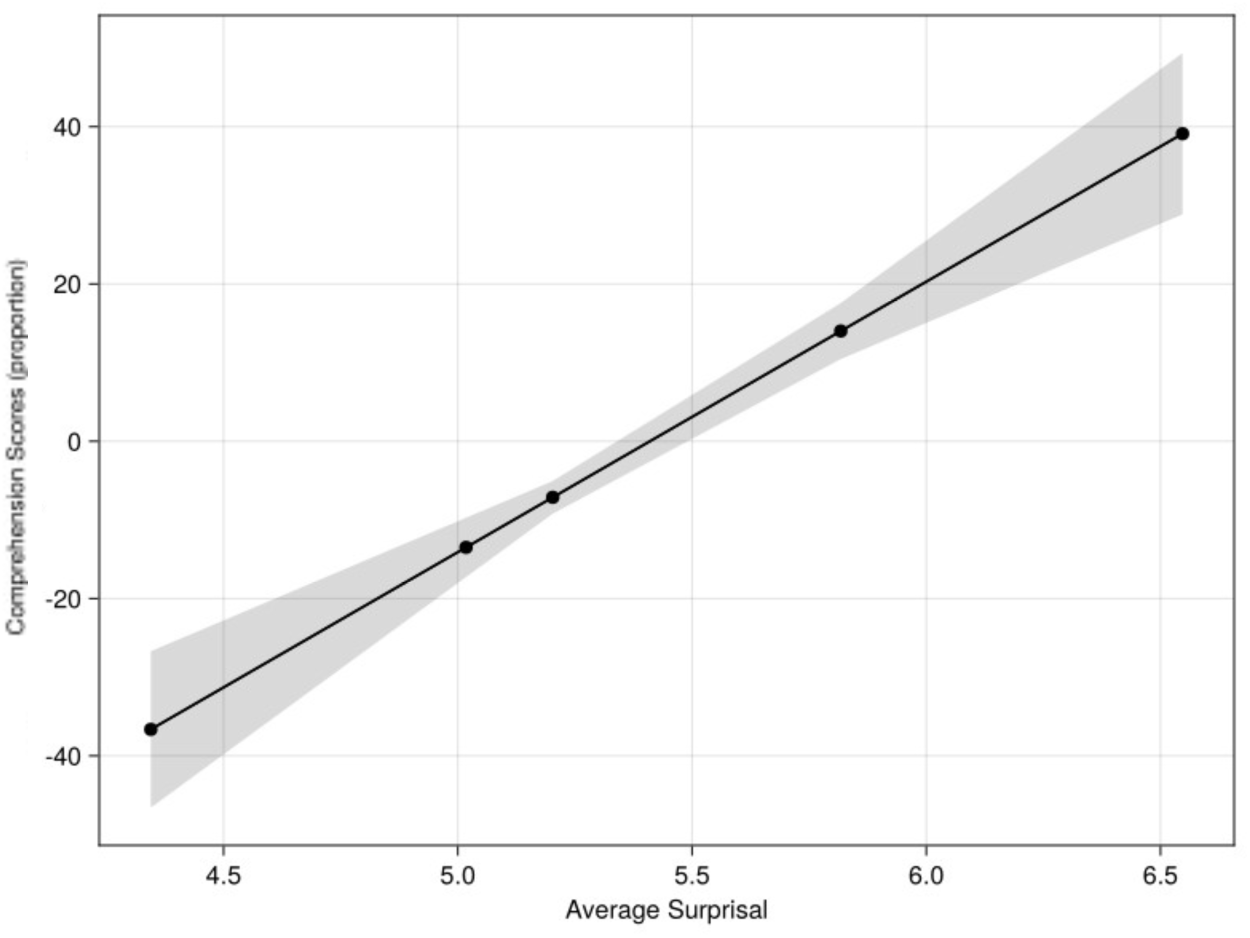
Main effect of average surprisal in CM4. The x-axis is average surprisal, whilst the y-axis represents fitted values of comprehension scores. The shaded areas represent 95% confidence intervals.

**Figure 11:**
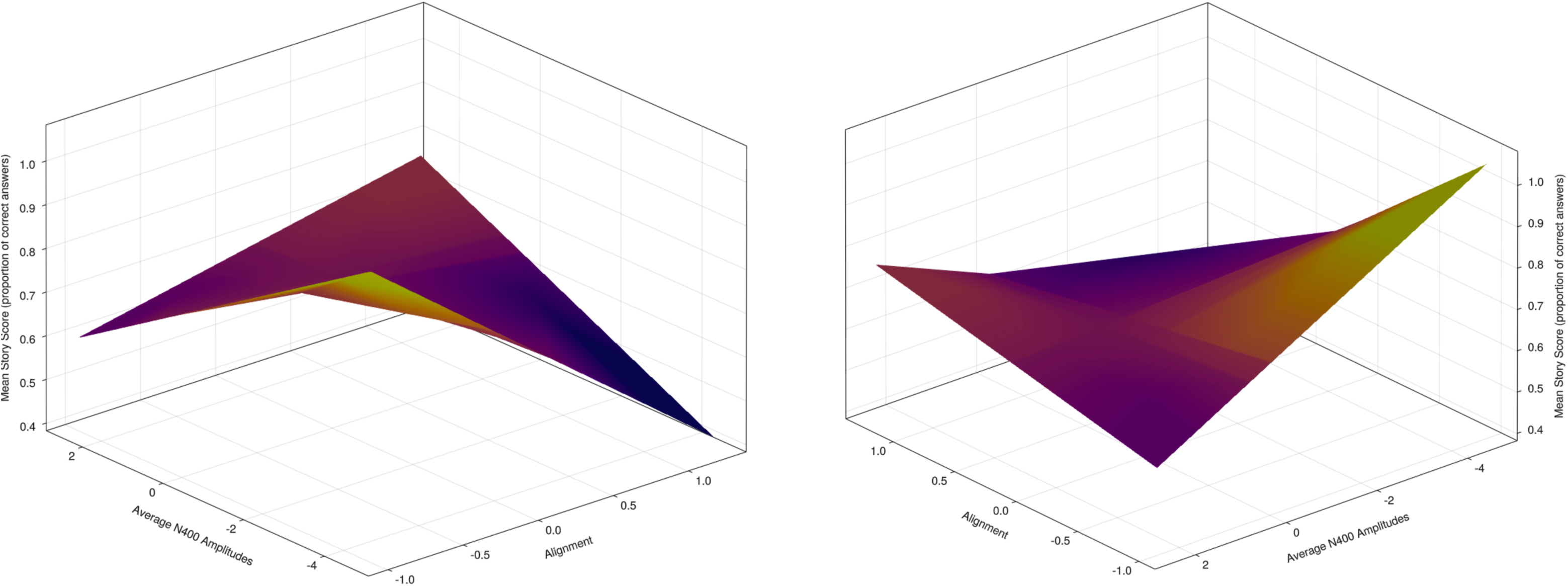
3D Surface Plot of CM1 highest order significant interaction effect between EEG amplitudes, alignment and depth of comprehension. Plots are coloured by depth of comprehension scores, where lighter colours represent better comprehension. The visualisation displays two isometric views rotated 90 degrees from each other.

#### CM2: Traditional N400 ROI & Surprisal

Main effects of both surprisal (Estimate = 23.46 Std. Error = 6.88, z = 3.39, *p* <.001) and surprisal^2^ (Estimate = −4.52, Std. Error = 1.29, z = −3.50, *p* <.001) were present in this model. Additionally, the interaction between surprisal and surprisal^2^ was also significant (Estimate = 0.29, Std. Error = 0.08, z = 3.61, *p* <.001 Figure 9 illustrates the effect of surprisal and demonstrates positive relationship between surprisal and comprehension, where higher surprisal is related to better comprehension. The significance of surprisal^2^ and its interaction with surprisal suggests the presence of a non-linear relationship.

#### CM3: Late Negativity ROI & EEG

No higher-order interaction effects were significant in this model. However, average EEG amplitude^2^ was an additional main effect (Estimate = −0.05, Std. Error = 0.03, z = −2.02, p = 0.04). Figure 12 visualises the non-significant interaction of average EEG amplitudes and alignment to demonstrate model effects when alignment is not a significant predictor of comprehension scores.

**Figure 12:**
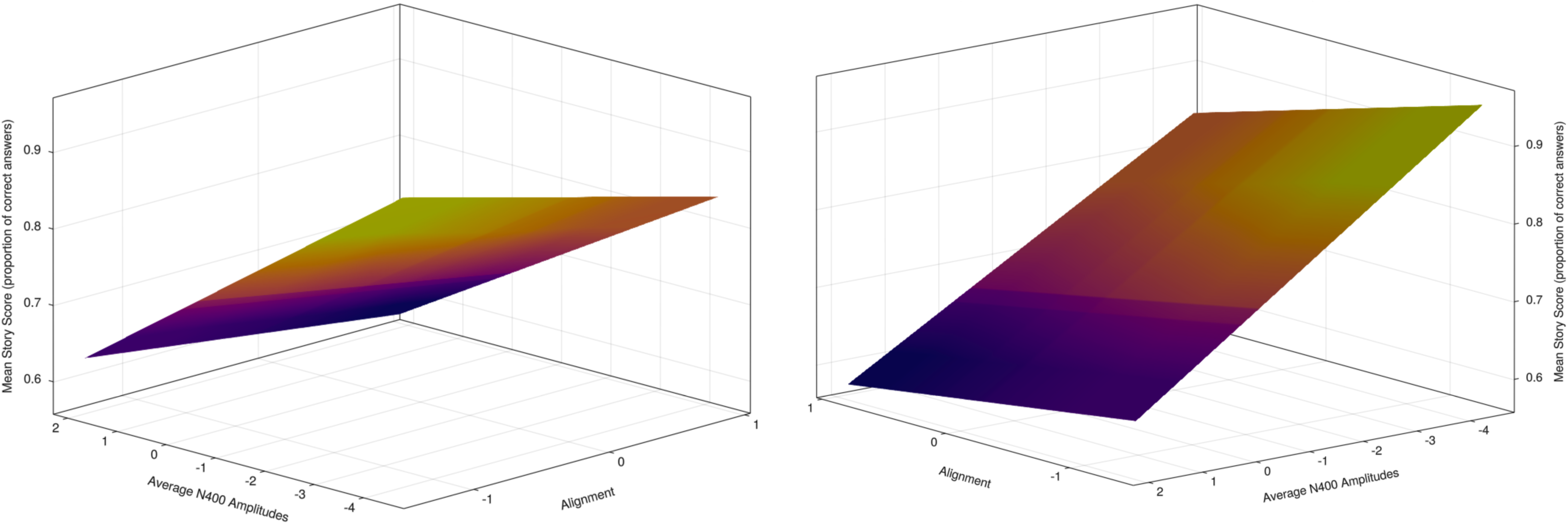
3D Surface Plot of CM3 non-significant highest order interaction effect between EEG amplitudes, alignment and depth of comprehension. Plots are coloured by depth of comprehension scores, where lighter colours represent better comprehension. The visualisation displays two isometric views rotated 90 degrees from each other.

#### CM4: Late Negativity ROI & Surprisal

Consistent with CM2, surprisal (Estimate = 25.2, Std. Error = 6.83, z = 3.69, *p* <.001), surprisal^2^ (Estimate = −4.87, Std. Error = 1.28, z = −3.81, *p* <.001) and surprisal and surprisal^2^ (Estimate = 0.31, Std. Error = 0.08, z = −1.60, p <.001) were significant predictors of comprehension scores. Figure 10 illustrates the effect of surprisal and demonstrates a positive relationship between surprisal and comprehension, where higher surprisal is related to better comprehension.

## 5 Discussion

The present study aimed to test the predictions outlined in the previous account of *Functional Unexpectedness* in language, with a specific focus on the role of contextual alignment. The framework proposed that higher alignment to local linguistic context would be associated with greater capacity to integrate unexpected information into existing schemas and therefore to access this new information during the comprehension task. To explore these relationships, the current study used two novel metrics to measure alignment and depth of comprehension. Four comprehension models were computed to explore the relationship between predictability (average EEG amplitudes or surprisal), alignment and comprehension depth. Three of these models revealed a main effect of predictability, with increasing depth of comprehension associated with increasing unexpectedness, as reflected by more negative EEG amplitudes or larger surprisal values. The traditional N400 EEG model (CM1) also demonstrated a role of alignment in predicting comprehension depth but, contrary to our hypothesis, revealed that lower alignment values were related to greater comprehension depth. Further, the comprehension models also demonstrated the presence of a non-linear relationship between predictability and comprehension depth, in support of the initial account, which outlined that a linear relationship may not best explain this interaction. In the following, we first outline these findings, followed by a discussion of the novel metrics and a consideration for how these results can be integrated into the *Functional Unexpectedness* account.

### 5.1 Evidence for Functional Unexpectedness

The two surprisal-based models (CM2 & CM4) revealed a functional role of surprisal, suggesting that unexpected stories were comprehended more effectively than less surprising stories. Although the initial account of Functional Unexpectedness suggested that this functionality would be based on contextual alignment at the individual and story level, this finding is in line with previous studies exploring the role of prediction errors in memory and learning. Outside of language processing, unexpectedness has been suggested to support the updating of internal predictive models, and these studies have demonstrated better memory performance (Haeuser & Kray, 2023; Hodapp & Rabovsky, 2021) and language acquisition (Lai et al., 2019) in the context of prediction errors during language processing. Therefore, the findings in the present study build on the foundation set by memory and learning studies and extend a functional role of unexpectedness to narrative comprehension. This is a novel finding in the context of naturalistic language comprehension and the measurement of comprehension depth. The presence of a significant quadratic and cubic (surprisal*surprisal^2^) predictor suggests that this relationship may not be best explained by a linear relationship alone. In the case of the quadratic predictor, a negative coefficient suggests an inverted U-shape relationship. This is in opposition to the specific U-shape predicted by functional unexpectedness, and as motivated by SLIMM. An inverted U instead suggests that a middle-ground level of surprisal is associated with the greatest comprehension. This pattern has been observed in attention research and labelled the ‘Goldilocks Effect’. Initially this effect was evident in explorations of infant auditory attention, where children are more likely to dedicate attention to events that are neither too simple (predictable) or too complex (unexpected) (Kidd et al., 2014; Kinney & Kagan, 1976)/ Exploring this concept in adults, Cao et al. (2025) showed that adults learnt novel verbs better after observing unexpected physical events (e.g., an object appearing to pass through solid wall) compared to expected events (e.g., an object stopped by the wall). Additionally, this ‘surprise-induced learning’ (Cao et al., 2025) exhibited a Goldilocks pattern, where learning initially increased in line with unexpected events but diminished when too many violations occurred. In the context of the present findings, this suggests a potential novelty threshold where information which is ‘too surprising’ may not be able to be integrated into existing schemas. Under the SLIMM framework, van Kesteren et al. (2012) suggested that encountering a novel event in a novel context will not lead to maximal learning as prior predictability distributions are less precise which may provide an explanation for the surprisal threshold evident in the surprisal models.

Complementing these surprisal models, the late negativity EEG model also demonstrated a functional role of unexpectedness, with more negative EEG amplitudes predicting better comprehension scores. Although this late negativity ROI falls outside of the traditional N400 window of 300-500ms we could consider that this is a shifted N400 effect, with a longer latency due to the processing of naturalistic language stimuli. In a study by Alday et al. (2017), in which participants listened to a 23 minute story, a similar shifted N400 effect was evident. Across several ERP analyses, the expected N400 frequency effect occurred around 700ms after word onset, rather than the traditional 400ms peak. Alday et al. (2017) noted that this was because word level effects in a naturalistic context reflect the sum of this context and additional complex interactions. Indeed, the processing of a continuous linguistic stream induces a cascade of brain activity and encompasses the processing of multiple linguistic levels, such as phonological processing, lexical access, semantic and syntactic integration (Sassenhagen, 2019). Neural activity for the current word will consist of a mixing of neural responses from words prior and upcoming, with word length also influencing this mixing (Alday et al., 2017; Sassenhagen, 2019). If we consider these results as evidence of a delayed N400 effect, then the present findings suggest that larger negativities in this time window are associated with better downstream comprehension. This finding would support our proposal of Functional Unexpectedness and sits alongside predictability conclusions from memory and learning rather than traditional language processing conclusions (i.e., larger N400 effects associated with less efficient comprehension).

### 5.2 An unexpected role of alignment

The traditional N400 ROI EEG model showed an interaction effect of alignment, EEG amplitudes and EEG amplitudes^2^, reflecting better comprehension for more negative EEG amplitudes for lower alignment values. Although this demonstrates a functional role of unexpectedness, the effect of alignment is in the opposite direction to the predictions Platt (2025). If the alignment metric was successful in capturing the strength of an individual’s existing predictive schemas, we may have been incorrect in our assumptions regarding how these schemas support comprehension outcomes. Our naturalistic stimuli were chosen to reflect language that individuals would experience outside of an experimental context. As such, they were sourced from short fictional stories, newspaper excerpts and popular science pieces. As a result, these stories were not designed to be difficult to comprehend or to present particularly challenging predictive conditions. Although this is a benefit of naturalistic stimuli, it may have produced conditions which did not require individuals to be highly aligned to the local context in order to fully understand the content. Under these conditions, individuals who were less aligned may have been experiencing more unexpected continuations, than those who were more aligned and their improved comprehension was a result of this functional relationship. This perspective could also be extended to precision-based accounts of comprehension, which suggest that model updating is based on the precision of the incoming signal and the strength of existing predictive models (Adams et al., 2013; Bornkessel-Schlesewsky & Schlesewsky, 2019; Feldman & Friston, 2010). We can assume that in the current experimental context signal precision is high and relatively consistent between participants, as story volume was chosen for the best listening experience and could be adjusted by the individual to best support hearing. Therefore, the difference between lower and higher aligned individuals is only in the strength of their existing predictive models tailored to each story. It may be that lower aligned individuals have less developed predictive models under certain story contexts and therefore are more likely to update these models following prediction errors. In comparison, higher aligned individuals would have stronger story-level predictive models and may be less likely to update their prediction models during the listening task. These proposed differences in model updating likelihood have been explored in relation to ageing (Moran et al., 2014) As the brain ages, a lifetime of experience has resulted in optimised prediction models which can minimise surprisal with minimal complexity. Moran et al. (2014) suggests that these optimised prediction models rely on top-down predictions and are less predisposed to short-term updating. In the context of the present results, higher aligned individuals may be processing unexpected stories with more optimised models (equivalent to an ‘older brain’; Moran et al. (2014) and are therefore less likely to integrate new information into these existing models for use during the comprehension ask. Similarly, Bornkessel-Schlesewsky et al. (2022) propose a comparable explanation for differences in short-term model updating across inter-individual differences with greater idea density, higher individual alpha frequency and steeper aperiodic 1/f slope associated with greater (‘younger brain’) model updating. These differences provide insights into the mechanisms underlying predictive model updating and the factors make adaptation more likely.

Additionally, our assumption that individuals who respond in line with surprisal values are better placed to utilise unexpected information to support comprehension relies on the belief that responding in an ‘average’ or expected way is the ideal. Although surprisal calculations take into consideration extended prior context and the statistical contingencies of language, these values may not reflect the ideal conditions for processing language. Rather than developing predictive models and associated schemas from a lifetime of unique experience, these models can only rely on learned associations held by their ‘attention heads’ (in the case of transformer models; Vaswani et al., 2017). Oh & Schuler (2023) provide evidence that larger transformer models, with more attention heads, are less predictive of human reading times, due to their greater capacity to learn statistical regularities, compared to humans. This finding suggests that larger LLMs may not apply predictions in a comparable way to human language processing. Findings from the present experiment, which suggest that predicting in line with LLM surprisal (higher alignment) was not associated with greater comprehension. Instead, lower alignment was associated with greater comprehension in response to prediction errors. This finding was justified by the suggestion that lower alignment indicated less precise schemas which were more likely to shift model updating to incoming stimuli. However, this outcome could also reflect that individuals who respond away from LLM surprisal, perhaps making predictions based on experiences which are not captured by the statistics of language, supports greater comprehension depth. Both of these perspectives provide insight into how individuals use the context available to them to support language comprehension and may indicate that further comparison of LLM surprisal and measures of language comprehension is required.

### 5.3 Further alignment considerations

To explore the relationship between predictability and language comprehension, the present study operationalised two novel metrics. Firstly, it used the established relationship between N400 amplitudes and surprisal to calculate an individual- and story-level alignment metric as a proxy for how well individuals were able to make contextually relevant predictions about upcoming linguistic stimuli. To produce these values, we computed an alignment model which predicted word-by-word N400 amplitudes, and which showed an interaction effect of surprisal and word frequency. Figure 5 demonstrates how the relationship between N400 amplitudes and surprisal changes for low frequency (i.e., more unexpected) words across the story duration, suggesting that as the story progresses low frequency words become less unexpected, as indexed by smaller (more positive) N400 amplitudes. This may reflect how individuals adjust predictions to local context as the story progresses and is in line with EEG research which suggests that N400 amplitudes can reflect an adaptation of predictions to local linguistic context (Hodapp & Rabovsky, 2021; Nieuwland & Van Berkum, 2006). Previous sentence-level findings indicate that frequency effects are strongest early on within a linguistic context (e.g., Payne et al., 2015; Alday et al., 2017). Alday et al. (2017) suggest that, as stories progress, relative frequency, as determined by the frequency of words within the current linguistic context, becomes a more accurate model to base predictions than corpus frequency (i.e., frequency of words within a global linguistic context). Although our surprisal metric is tailored to local context, it still relies on the global frequency of words, which may suggest that our alignment metric could be improved by considering relative frequency as well.

However, evidence of a frequency effect in this alignment relationship (N400 ∼ surprisal) suggests that our alignment metric also has the capacity to take this effect into account in our subsequent comprehension models. Effects of word frequency on the N400 component have been shown to be context-dependent and modulated by story position. Evidence of this effect in the alignment models provides an indicator of effectiveness of the alignment metric. Encompassing the frequency effect is essential as findings from eye-tracking literature suggest that substantial variation in processing times, as evidence by reading and fixation times, can be explained in reference to low level features such as word frequency (for a review see, Rayner, 1998). Higher frequency words have been shown to have shorter fixation durations and reading times than low frequency words and this is often considered a reflection of decreased processing effort (e.g., Ashby et al., 2005; Calvo & Meseguer, 2002; Huck et al., 2017; Kennedy et al., 2013). As our model indicates a different relationship between N400 amplitudes and surprisal by word frequency, it suggests that our alignment metric had the capacity to account for the influence of word frequency on language comprehension at a word-by-word level for each individual per story. Further supporting the proposal that the relationship between N400 amplitudes and surprisal within the alignment metric can capture aspects of how the linguistic context is processed.

### 5.4 Comprehension depth

A series of comprehension questions were designed to reflect an individual’s understanding of each story. More specially, the aim of these questions was to require listeners to piece together information from the stories which was not directly stated. For example, instead of only asking a character’s name or age, we also asked about the author’s opinion of this person based only on the way they were written about or about an underlying message within the text. As the comprehension questions rely on a contextual understanding of stories, it was assumed that this type of information use would be more comparable to everyday communication, where individuals often draw inferences based on provided information. The average comprehension depth score across all genres was 70%, suggesting that the questions designed did not reach a ceiling effect but that individuals were still answering above chance. Additionally, the range of comprehension scores was 50-86.1% indicating that there was a reasonable amount of variability in scores between individuals, allowing us to explore how these comprehension differences may have arisen.

It was important to evaluate the suitability of the depth of comprehension metric within this context as it differs from traditional comprehension metrics in language comprehension studies. One opportunity for comparison and evaluation is with the control predictors also included in this analysis. Overall, the science genre stories had the lowest average comprehension score (67.4%), compared with newspaper (69.7%) and fiction (72.9%) stories. These stories also had (equal) highest idea density, a measure in which higher idea density (i.e., increased propositional density) has been associated with a decrease in language comprehension efficiency (Harrag et al., 2025). It is thought that the increased complexity of these texts requires more processing effort than lower ID text, therefore influencing overall comprehension. Additionally, the science stories also had the longest average word duration, which has also been associated with (Gerth & Festman, 2021). In the opposite case, the higher scoring fiction stories had the lowest word frequency (Dambacher et al., 2006), word duration and shortest sentence length. These converging results provide evidence for the effectiveness of the depth of comprehension measure.

### 5.5 Functional unexpectedness: Future directions

Although our analysis approach has provided some support for our predictions regarding a functional role of unexpectedness in language comprehension and that this functionality may differ by individual-level story alignment, there are still some areas for further development and consideration. Most crucially, there is a further need to understand the role of alignment in utilising unexpected information. In light of the current findings, the account of Functional Unexpectedness needs to be expanded to consider multiple predictive contexts. For example, we could consider whether higher alignment is required to elicit functional unexpectedness in a less general context such as testing non-experts versus experts on a niche research topic (e.g., graded Harry Potter knowledge; Troyer & Kutas, 2020). This will require future research to explore functional unexpectedness and measure alignment across a range of linguistic contexts to indicate whether the role of alignment differs. This would also provide the opportunity for more unexpected information to be presented without jeopardizing the naturalistic nature of stimuli while still expanding the range of predictability. Further, it may be necessary to consider whether individuals responding at a baseline level (i.e., in line with GPT-2 surprisal estimates) is ideal for increased comprehension outcomes or whether alternative modes of surprisal calculations may provide a more human representative baseline (e.g., via smaller language models as suggested in Oh & Schuler, 2023). Our findings suggest that at least in this context, individuals who were less aligned with surprisal at a story level had better comprehension scores for larger N400 amplitudes, suggesting that our assumptions around ideal predictive conditions may not have been correct. If the relationship between lower alignment and better comprehension can be replicated in additional contexts, this finding may provide insight into the mechanisms underlying model updating and the role of schema strength in this process.

Finally, there is still an opportunity for the comprehension analysis to be more fine-grained, rather than including primarily story-level averaged predictors. Prior research has suggested that individuals adapt their predictions across story comprehension and that this shapes the predictability of information. We demonstrate this change in the alignment models, which indicate that the relationship between N400 amplitudes and surprisal changes as the story progress and that this is especially prominent for low frequency words. Fortunately, our alignment metric is dictated by a word-level alignment model and by design, is included to incorporate across story change. However, future analyses can further separate story-level predictors within the comprehension analysis. This could include sectioning the naturalistic stories at relatively natural breaks to allow for section-level averaging. A sectioned approach would allow for alignment to be measured for each section and key predictors (e.g., surprisal and N400 amplitudes) can be averaged at this level to further reflect how predictions change as more linguistic information becomes available. Further, comprehension questions could be added within these breaks to measure how story understanding changes in line with alignment. This approach could add an additional level of naturalistic experience as naturalistic linguistic stimuli are often accompanied by natural breaks in discourse, such as book chapters, advertising breaks in television and naturalistic conversations which operate between periods of speaking, listening and thinking. To further understand the downstream effects of unexpectedness future studies could also include a measure of memory retention at a later time. This could provide further insight into influence of timing on the relationship between predictability and behavioural outcomes such as depth of comprehension and memory.

## 6 Conclusion

In language processing, individuals are tasked with processing unexpected linguistic inputs. Evidence from language studies which focus on the real-time processing of language suggest that unexpected words are harder to process and that this negatively impacts language comprehension. The present study has demonstrated that when measuring comprehension depth (i.e., a contextual understanding of naturalistic language stimuli), comprehension is improved for unexpected stories, rather than reduced. Additionally, our findings suggest that there may be further complexity in this functional relationship which may be better explored using quadratic or cubic predictors. Our novel alignment metric also demonstrated that individuals who are less aligned to local context (at the individual and story level) can better utilise unexpected information than higher aligned individuals. This suggests that an individual’s internal predictive conditions influence the relationship between predictability and comprehension. These conclusions provided experimental evidence for predictions outlined in the account of *Functional Unexpectedness* and suggest that further consideration for the how individuals utilise existing schemas during language comprehension is required. Additionally, although the prediction that the relationship between predictability and comprehension depth would be modulated by alignment, the directionality of this effect was not as predicted. More broadly, these findings have implications for how the role of unexpectedness is viewed in the processing of language and suggests that there are opportunities for this characteristic to be utilised to improve comprehension in naturalistic communication.

## Supplementary Material A: Naturalistic story stimuli examples

### F02 – The new food

I see from the current columns of the daily press that “Professor Plumb, of the University of Chicago, has just invented a highly concentrated form of food. All the essential nutritive elements are put together in the form of pellets, each of which contains from one to two hundred times as much nourishment as an ounce of an ordinary article of diet. These pellets, diluted with water, will form all that is necessary to support life. The professor looks forward confidently to revolutionizing the present food system. “Now this kind of thing may be all very well in its way, but it is going to have its drawbacks as well. In the bright future anticipated by Professor Plumb, we can easily imagine such incidents as the following:

The smiling family were gathered round the hospitable board. The table was plenteously laid with a soup-plate in front of each beaming child, a bucket of hot water before the radiant mother, and at the head of the board the Christmas dinner of the happy home, warmly covered by a thimble and resting on a poker chip. The expectant whispers of the little ones were hushed as the father, rising from his chair, lifted the thimble and disclosed a small pill of concentrated nourishment on the chip before him. Christmas turkey, cranberry sauce, plum pudding, mince pie--it was all there, all jammed into that little pill and only waiting to expand. Then the father with deep reverence, and a devout eye alternating between the pill and heaven, lifted his voice in a benediction. At this moment there was an agonized cry from the mother.

“Oh, Henry, quick! Baby has snatched the pill!” It was too true. Dear little Gustavus Adolphus, the golden-haired baby boy, had grabbed the whole Christmas dinner off the poker chip and bolted it. Three hundred and fifty pounds of concentrated nourishment passed down the oesophagus of the unthinking child. “Clap him on the back!” cried the distracted mother. “Give him water!”

The idea was fatal. The water striking the pill caused it to expand. There was a dull rumbling sound and then, with an awful bang, Gustavus Adolphus exploded into fragments! And when they gathered the little corpse together, the baby lips were parted in a lingering smile that could only be worn by a child who had eaten thirteen Christmas dinners.

### N02 – Colour, spectators to greet 2000 ‘Tour’

Planning is well underway for Strathalbyn and Goolwa’s involvement in the Jacob’s Creek Tour Down Under Cycle Race on January 19. Both towns are making sure colour and excitement will be prominent features for the race which will have international TV coverage. Stage 2 starts at North Adelaide through the Adelaide Hills and into Strathalbyn for the sprint in Dawson Street and then into Goolwa for the race finish in Cadell Street. The towns will welcome the cyclists with their own decoration themes.

Strathalbyn will greet the cyclists with a large painted mural on the hillside at Castle Hill and a town entrance rural theme display leading into a tartan painted road and hundreds of coloured balloons through Dawson Street for the TAB Sprint. Goolwa’s theme will be river and paddle steamers entrance statement across Cadell Street from Moore Streets and colourful flags and sails to the finish line by the Goolwa Hotel.

Businesses and householders are encouraged to decorate their premises along the route in the tour’s colours of Red. Blue and Yellow to be judged by KESAB on the day. There are awards for the Best Decorated Business and Best Decorated House on route in each town and a special award for the Best Dressed Town overall. Entry forms for the Best Dressed Business and House competitions are available from Strathalbyn Tourism Information Centre and at Goolwa from Alexandrina Council Offices. Closing date for entries is Friday, January 14.

Roving entertainers, a Jazz Band, special race events, Sky Dive and a Micro Lite Fly By are just some of the events planned for January 19th 2000 when Goolwa hosts the finish of Stage 2 of the Jacob’s Creek Tour Down Under Cycle Race. Up to 100 of the world’s best cyclists will be taking part in this exciting race which will be enjoyed by an international television audience. The six stage race takes place in South Australia from January 18-23, 2000 and promises to be an even bigger event than the first ‘Tour Down Under’ earlier this year.

Pre-race street activities at Goolwa include the SA Veterans & Ladies Cycling Classic Race, Speed Skating, and a Mountain Bike Stunt Team in the children’s park behind Cadell Street where there will also be sponsors’ displays, food and drink stalls, children’s mini fair and face painting. Stuart O’Grady has autographed three bicycle helmets and Tour Down Under T-Shirts for prizes in a competition for under fifteen year olds to be judged on the spot on the day for the best banner or flag brought to cheer the cyclists as they arrive in Goolwa.

### S06 – New research sheds light on how dogs became dogs

At first blush, the emergence of man’s best friend is pretty straightforward. The first dogs descended from wolves in Europe about 14,000 years ago. Then humans domesticated those proto-dogs until the eventual animal known as a “dog” had many of the traits we associated with the animal today. That much of the evolutionary history of the modern dog has been clearly understood. But further research suggests that the European dog is not the ancestor of all our dogs; instead, every modern Western dog hails from a Southeast Asian progenitor lineage. Why? Why did some upstart Southeast Asian lineage triumph, even in Europe, instead of the endemic European one? Turns out, it might have to do with your pet dog’s affinity for Cheetos.

According to research conducted by Ben Sacks from the University of California at Davis and his colleagues, the Southeast Asian dogs prospered because, after they were brought south of the Yangtze River some 6,000 years ago, the dogs were isolated from their wolf forebears. Without that proximity, the Southeast Asian dogs could no longer interbreed with wolves, and thus followed their own evolutionary path. In contrast, northern Asian and European dogs still had contact with, and interbred with, the native wolf populations. Put more clearly, if dogs and wolves interbreed, as they did in Europe, they ended up in an evolutionary cul-de-sac. Isolated from one another, traits that benefited the newly emergent dog lineage flourished.

Another slice of data, published in January in Nature by Erik Axelsson of Uppsala University, suggests that one of the main differences between dogs and wolves is their ability to digest starch. As dogs continued their co-existence with humans - who were, at the same time, mastering agriculture and switching to a grain-based diet - those individuals who could eat starch would be better suited to a domesticated lifestyle than those who had to constantly hunt. Sacks and Axelsson disagree on when that switch took place: Axelsson says that it happened before humans switched from a hunter-gatherer lifestyle to a farming one; Sacks says that the mutation occurred once rice cultivation in Southeast Asia was well underway.

Further work will be required to pinpoint just when the modern dog eventually emerged and to clarify where other canids, such as dingoes, fit into this evolutionary picture. In the interim, sit back and marvel at the process that eventually produced the donut-stealing canine scamps we know and love today.

## Supplementary Material B: Model summaries

**Table SMB-1:**
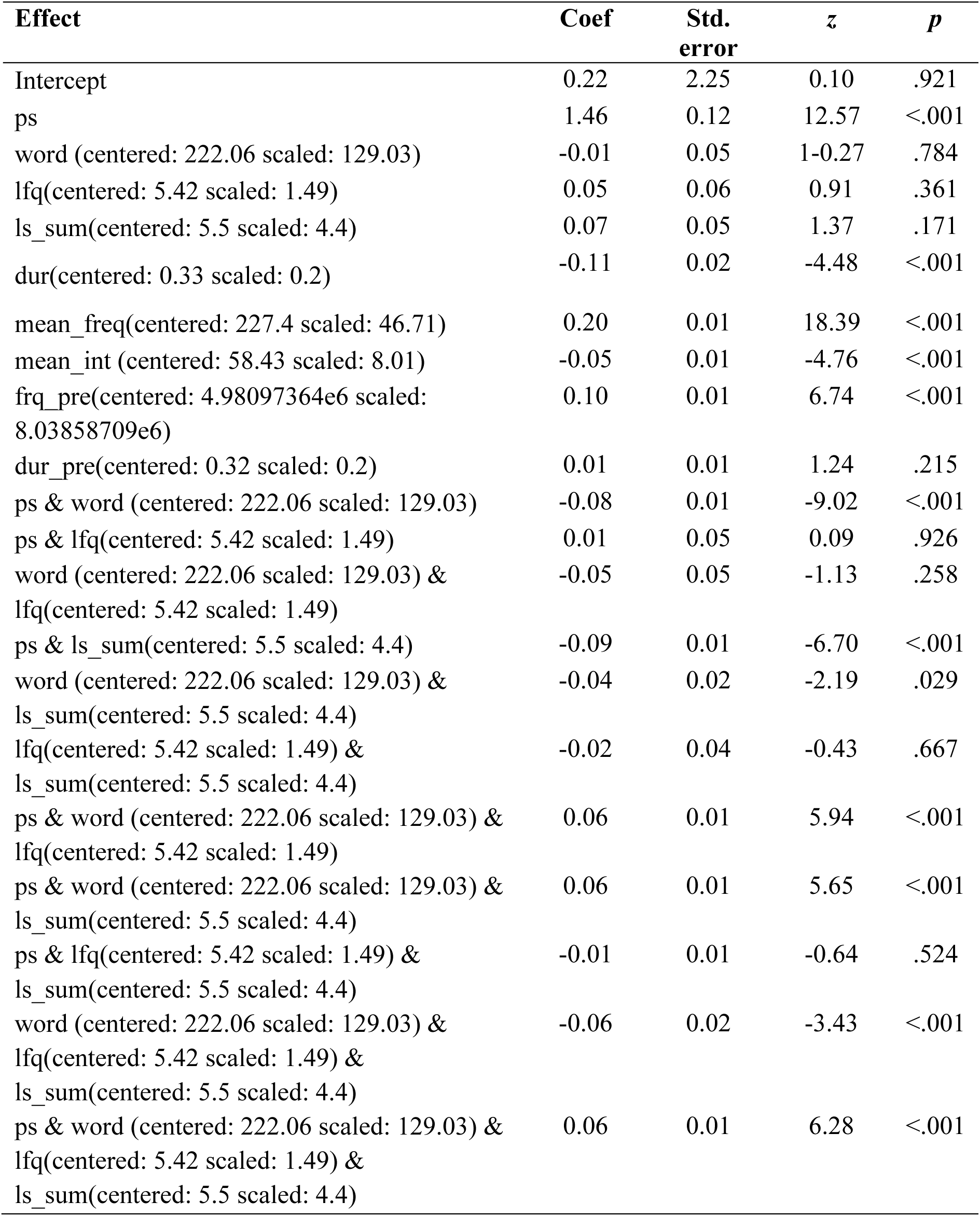
Model output for AM1.

| Effect | Coef | Std.<br>error | z | p |
| --- | --- | --- | --- | --- |
| Intercept | 0.22 | 2.25 | 0.10 | .921 |
| ps | 1.46 | 0.12 | 12.57 | <.001 |
| word (centered: 222.06 scaled: 129.03) | -0.01 | 0.05 | 1-0.27 | .784 |
| lfq(centered: 5.42 scaled: 1.49) | 0.05 | 0.06 | 0.91 | .361 |
| ls_sum(centered: 5.5 scaled: 4.4) | 0.07 | 0.05 | 1.37 | .171 |
| dur(centered: 0.33 scaled: 0.2) | -0.11 | 0.02 | -4.48 | <.001 |
| mean_freq(centered: 227.4 scaled: 46.71) | 0.20 | 0.01 | 18.39 | <.001 |
| mean_int (centered: 58.43 scaled: 8.01) | -0.05 | 0.01 | -4.76 | <.001 |
| frq_pre(centered: 4.98097364e6 scaled: 8.03858709e6) | 0.10 | 0.01 | 6.74 | <.001 |
| dur_pre(centered: 0.32 scaled: 0.2) | 0.01 | 0.01 | 1.24 | .215 |
| ps & word (centered: 222.06 scaled: 129.03) | -0.08 | 0.01 | -9.02 | <.001 |
| ps & lfq(centered: 5.42 scaled: 1.49) | 0.01 | 0.05 | 0.09 | .926 |
| word (centered: 222.06 scaled: 129.03) & lfq(centered: 5.42 scaled: 1.49) | -0.05 | 0.05 | -1.13 | .258 |
| ps & ls_sum(centered: 5.5 scaled: 4.4) | -0.09 | 0.01 | -6.70 | <.001 |
| word (centered: 222.06 scaled: 129.03) & ls_sum(centered: 5.5 scaled: 4.4) | -0.04 | 0.02 | -2.19 | .029 |
| lfq(centered: 5.42 scaled: 1.49) & ls_sum(centered: 5.5 scaled: 4.4) | -0.02 | 0.04 | -0.43 | .667 |
| ps & word (centered: 222.06 scaled: 129.03) & lfq(centered: 5.42 scaled: 1.49) | 0.06 | 0.01 | 5.94 | <.001 |
| ps & word (centered: 222.06 scaled: 129.03) & ls_sum(centered: 5.5 scaled: 4.4) | 0.06 | 0.01 | 5.65 | <.001 |
| ps & lfq(centered: 5.42 scaled: 1.49) & ls_sum(centered: 5.5 scaled: 4.4) | -0.01 | 0.01 | -0.64 | .524 |
| word (centered: 222.06 scaled: 129.03) & lfq(centered: 5.42 scaled: 1.49) & ls_sum(centered: 5.5 scaled: 4.4) | -0.06 | 0.02 | -3.43 | <.001 |
| ps & word (centered: 222.06 scaled: 129.03) & lfq(centered: 5.42 scaled: 1.49) & ls_sum(centered: 5.5 scaled: 4.4) | 0.06 | 0.01 | 6.28 | <.001 |

**Table SMB-2:** Model outputs for AM2.

| Effect | Coef | Std.<br>error | z | p |
| --- | --- | --- | --- | --- |
| Intercept | -0.10 | 0.05 | -1.93 | .053 |
| ps | 0.06 | 0.10 | 0.62 | .532 |
| word (centered: 222.06 scaled: 129.03) | -0.01 | 0.04 | -0.10 | .918 |
| lfq(centered: 5.42 scaled: 1.49) | 0.01 | 0.04 | 0.16 | .873 |
| ls_sum(centered: 5.5 scaled: 4.4) | -0.10 | 0.04 | -2.49 | .013 |
| dur(centered: 0.33 scaled: 0.2) | -0.01 | 0.01 | -0.51 | .6121 |
| mean_freq(centered: 227.4 scaled: 46.71) | 0.06 | 0.01 | 10.68 | <.001 |
| mean_int (centered: 58.43 scaled: 8.01) | -0.06 | 0.01 | -10.02 | <.001 |
| frq_pre(centered: 4.98097364e6 scaled:<br>8.03858521e6) | 0.01 | 0.01 | 1.36 | .175 |
| dur_pre(centered: 0.32 scaled: 0.2) | 0.10 | 0.01 | 15.57 | <.001 |
| ps & word (centered: 222.06 scaled: 129.03) | -0.07 | 0.01 | -12.73 | <.001 |
| ps & lfq(centered: 5.42 scaled: 1.49) | -0.05 | 0.04 | -1.36 | .175 |
| word (centered: 222.06 scaled: 129.03) & lfq<br>(centered: 5.42 scaled: 1.49) | 0.09 | 0.03 | 2.72 | .006 |
| ps & ls_sum(centered: 5.5 scaled: 4.4) | -0.14 | 0.01 | -20.06 | <.001 |
| word (centered: 222.06 scaled: 129.03) &<br>ls_sum(centered: 5.5 scaled: 4.4) | 0.05 | 0.01 | 5.34 | <.001 |
| lfq(centered: 5.42 scaled: 1.49) &<br>ls_sum(centered: 5.5 scaled: 4.4) | 0.04 | 0.03 | 1.07 | .287 |
| ps & word (centered: 222.06 scaled: 129.03) &<br>lfq(centered: 5.42 scaled: 1.49) | 0.05 | 0.01 | 8.46 | <.001 |
| ps & word (centered: 222.06 scaled: 129.03) &<br>ls_sum(centered: 5.5 scaled: 4.4) | 0.05 | 0.01 | 8.69 | <.001 |
| ps & lfq(centered: 5.42 scaled: 1.49) &<br>ls_sum(centered: 5.5 scaled: 4.4) | 0.01 | 0.01 | 0.09 | .931 |
| word (centered: 222.06 scaled: 129.03) &<br>lfq(centered: 5.42 scaled: 1.49) &<br>ls_sum(centered: 5.5 scaled: 4.4) | -0.10 | 0.01 | -9.61 | <.001 |
| ps & word (centered: 222.06 scaled: 129.03) &<br>lfq(centered: 5.42 scaled: 1.49) &<br>ls_sum(centered: 5.5 scaled: 4.4) | 0.06 | 0.01 | 11.80 | <.001 |

**Table SMB-3:**
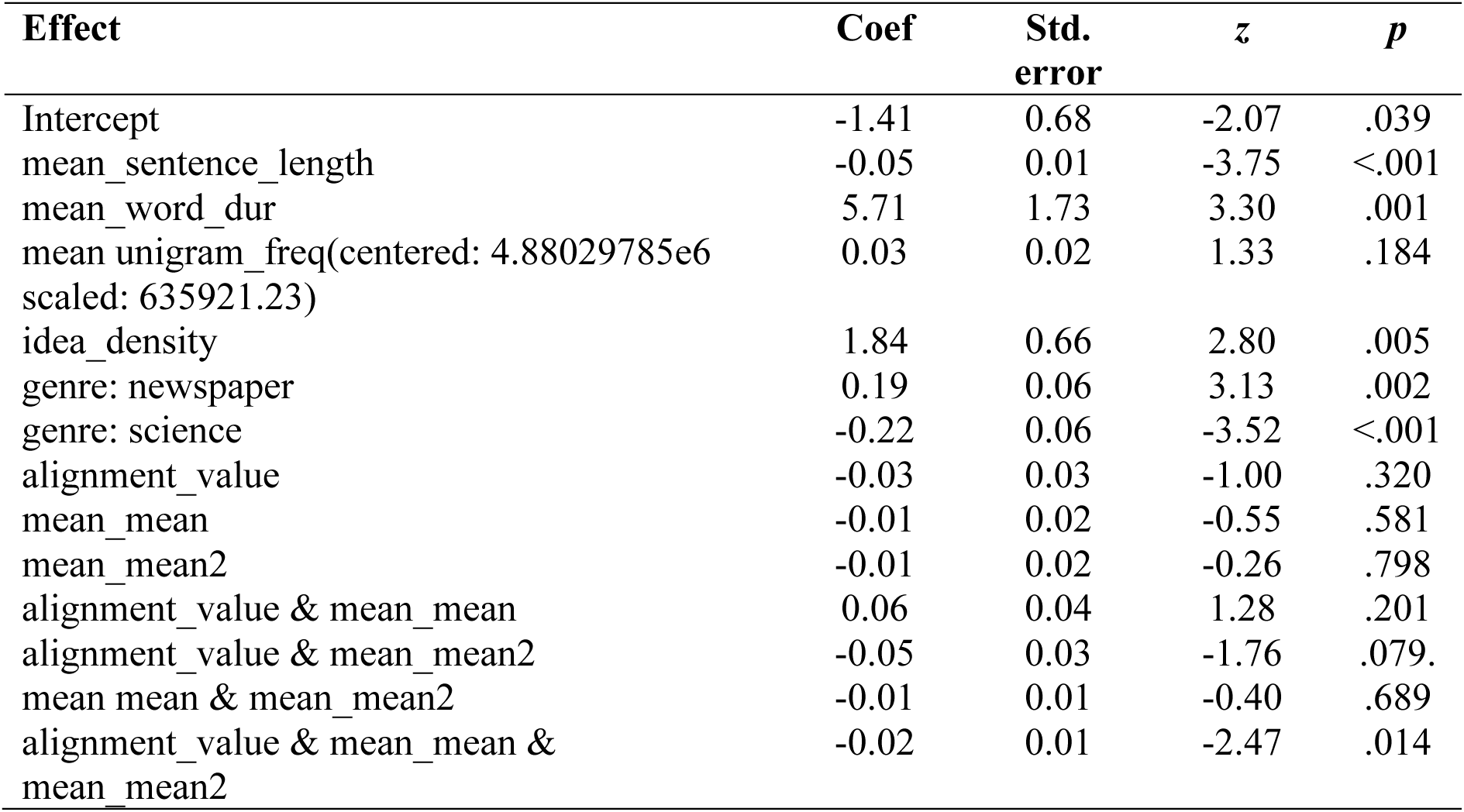
Model outputs for CM1.

| Effect | Coef | Std.<br>error | z | p |
| --- | --- | --- | --- | --- |
| Intercept | -1.41 | 0.68 | -2.07 | .039 |
| mean_sentence_length | -0.05 | 0.01 | -3.75 | <.001 |
| mean_word_dur | 5.71 | 1.73 | 3.30 | .001 |
| mean unigram_freq(centered: 4.88029785e6<br>scaled: 635921.23) | 0.03 | 0.02 | 1.33 | .184 |
| idea_density | 1.84 | 0.66 | 2.80 | .005 |
| genre: newspaper | 0.19 | 0.06 | 3.13 | .002 |
| genre: science | -0.22 | 0.06 | -3.52 | <.001 |
| alignment_value | -0.03 | 0.03 | -1.00 | .320 |
| mean_mean | -0.01 | 0.02 | -0.55 | .581 |
| mean_mean2 | -0.01 | 0.02 | -0.26 | .798 |
| alignment_value & mean_mean | 0.06 | 0.04 | 1.28 | .201 |
| alignment_value & mean_mean2 | -0.05 | 0.03 | -1.76 | .079. |
| mean_mean & mean_mean2 | -0.01 | 0.01 | -0.40 | .689 |
| alignment_value & mean_mean &<br>mean_mean2 | -0.02 | 0.01 | -2.47 | .014 |

**Table SMB-4:** Model outputs for CM2.

| Effect | Coef | Std.<br>error | z | p |
| --- | --- | --- | --- | --- |
| Intercept | -43.30 | 12.43 | -3.48 | <.001 |
| mean_sentence_length | -0.10 | 0.01 | -7.5 | <.001 |
| mean_word_dur | 9.20 | 1.60 | 5.74 | <.001 |
| mean unigram_freq(centered: 4.88029785e6<br>scaled: 635921.23) | 0.13 | 0.02 | 4.34 | <.001 |
| idea_density | 4.45 | 0.73 | 6.34 | <.001 |
| genre: newspaper | 0.32 | 0.05 | 6.80 | <.001 |
| genre: science | -0.29 | 0.06 | -5.32 | <.001 |
| alignment_value | 1.04 | 14.82 | 0.07 | .944 |
| ls_sum_mean_fixed | 23.36 | 6.88 | 3.39 | <.001 |
| ls_sum_mean2_fixed | -4.42 | 1.29 | -3.50 | <.001 |
| alignment_value & ls_sum_mean_fixed | -0.30 | 8.39 | -0.04 | .971 |
| alignment_value & ls_sum_mean2_fixed | 0.01 | 1.57 | 0.00 | .996 |
| ls_sum_mean_fixed & ls_sum_mean2_fixed | 0.29 | 0.08 | 3.61 | <.001 |
| alignment_value & ls_sum_mean_fixed &<br>ls_sum_mean2_fixed | 0.01 | 0.10 | 0.02 | .982 |

**Table SMB-5:** Model outputs for CM3.

| Effect | Coef | Std.<br>error | z | p |
| --- | --- | --- | --- | --- |
| Intercept | -1.39 | 0.69 | -2.02 | .044 |
| mean_sentence_length | -0.05 | 0.01 | -3.61 | <.001 |
| mean_word_dur | 5.70 | 1.78 | 3.24 | .001 |
| mean unigram_freq(centered: 4.88029785e6<br>scaled: 635921.23) | 0.03 | 0.03 | 1.19 | .233 |
| idea_density | 1.79 | 0.66 | 2.70 | .007 |
| genre: newspaper | 0.19 | 0.06 | 3.03 | .002 |
| genre: science | -0.21 | 0.06 | -3.41 | <.001 |
| alignment_value | -0.02 | 0.03 | -0.47 | .636 |
| mean_mean | -0.04 | 0.02 | -1.75 | .080 |
| mean_mean2 | -0.05 | 0.03 | -2.02 | .043 |
| alignment_value & mean_mean | 0.01 | 0.05 | 0.16 | .872 |
| alignment_value & mean_mean2 | -0.06 | 0.05 | -1.21 | .227 |
| mean_mean & mean_mean2 | -0.01 | 0.01 | -1.96 | .050 |
| alignment_value & mean_mean &<br>mean_mean2 | -0.02 | 0.01 | -1.70 | .089 |

**Table SMB-6:** Model outputs for CM4.

| Effect | Coef | Std.<br>error | z | p |
| --- | --- | --- | --- | --- |
| Intercept | -46.49 | 12.34 | -3.77 | <.001 |
| mean_sentence_length | -0.10 | 0.01 | -7.51 | <.001 |
| mean_word_dur | 9.05 | 1.58 | 5.71 | <.001 |
| mean unigram_freq(centered: 4.88029785e6<br>scaled: 635921.23) | 0.13 | 0.03 | 4.35 | <.001 |
| idea_density | 4.63 | 0.72 | 6.41 | <.001 |
| genre: newspaper | 0.32 | 0.05 | 6.83 | <.001 |
| genre: science | -0.29 | 0.05 | -5.32 | <.001 |
| alignment_value | -23.69 | 16.49 | -1.44 | .151 |
| ls_sum_mean_fixed | 25.21 | 6.83 | 3.69 | <.001 |
| ls_sum_mean2_fixed | -4.87 | 1.28 | -3.81 | <.001 |
| alignment_value & ls_sum_mean_fixed | 14.17 | 9.36 | 1.51 | .130 |
| alignment_value & ls_sum_mean2_fixed | -2.81 | 1.76 | -1.60 | .110 |
| ls_sum_mean_fixed & ls_sum_mean2_fixed | 0.31 | 0.08 | 3.92 | <.001 |
| alignment_value & ls_sum_mean_fixed &<br>ls_sum_mean2_fixed | 0.18 | 0.11 | 1.68 | .093 |

## Notes

### Competing Interest Statement

The authors have declared no competing interest.

